# Cutaneous lupus features specialized stromal niches and altered retroelement expression

**DOI:** 10.1101/2025.02.01.636053

**Authors:** Jeff R. Gehlhausen, Yong Kong, Emily Baker, Sarika Ramachandran, Fotios Koumpouras, Christine J. Ko, Matthew Vesely, Alicia J. Little, William Damsky, Brett King, Akiko Iwasaki

**Author notes:** COI: A.I. co-founded RIGImmune, Xanadu Bio and PanV; and is a member of the Board of Directors of Roche Holding Ltd and Genentech. J.R.G has received research funding support from Bristol Meyers Squibb.

## Abstract

Cutaneous Lupus is an inflammatory skin disease causing highly morbid inflamed skin and hair loss. In order to investigate the pathophysiology of cutaneous lupus, we performed single-cell RNA and spatial sequencing of lesional and non-lesional cutaneous lupus skin compared to healthy controls. Pathway enrichment analyses of lesional keratinocytes revealed elevated responses to type I interferon, type II interferon, tumor necrosis factor, and apoptotic signaling. Detailed clustering demonstrated unique fibroblasts specific to lupus skin with likely roles in inflammatory cell recruitment and fibrosis. We also evaluated the association of retroelement expression with type I interferons in the skin. We observed increased retroelement expression which correlated with interferon-stimulated genes across multiple cell types. Moreover, we saw elevated expression of genes involved in RIG-I and cGAS-STING pathways, which transduce elevated nucleic acid signals. Treatment of active cutaneous lupus with Anifrolumab reduced RIG-I and cGAS-STING pathways in addition to the most abundant retroelement family, L2b. Our studies better define type I interferon-mediated immunopathology in cutaneous lupus and identify an association between retroelement expression and interferon signatures in cutaneous lupus.

## Introduction

Cutaneous Lupus Erythematosus (CLE) is an autoimmune skin disease affecting over 200,000 individuals in the United States.(1) While CLE is often seen in association with Systemic Lupus Erythematosus (SLE), it is also seen in many patients without SLE. CLE, in its most common Discoid subtype, is characterized by inflamed, scarring plaques typically on the head and neck that can be highly disfiguring. To date, there are no FDA-approved medications specific for the treatment of CLE and half of cases are refractory to initial measures that include topical corticosteroids and antimalarials.(2)

Though incompletely understood, the pathophysiology of CLE involves type I interferon-mediated effects on one or more critical cell types in the microenvironment.(3–5) Histopathologically, the accumulation of plasmacytoid dendritic cells (pDCs) is specific for CLE and is associated with the elevated type I interferon (IFN-I) signatures. pDCs are unique in their ability to produce massive amounts of IFN-α upon stimulation and have long been hypothesized as the major producer of IFN-I in CLE.(5) Recent studies, however, suggest that pDCs may not be the major producers of type IFN-I in the CLE microenvironment.(6,7) Nonetheless, pDCs appear to have a central role in CLE given that multiple new pDC-specific antibodies have demonstrated encouraging results in clinical trials.(8,9) Moreover, the IFNAR-blocking antibody Anifrolumab is also showing promising results in the treatment of CLE, further underscoring the central role of the IFN-I pathway in CLE.(10) In addition to IFN-I, tumor necrosis factor (TNF), interferon gamma, and Fas-Fas ligand signaling have been implicated as other pathways potentially involved in CLE pathogenesis.(4)

Downstream of IFN-I, the major drivers of immunopathology in CLE also remain an area of intense investigation. Prior work has shown IFN-Is can lower the threshold for apoptosis in keratinocytes – which may be a critical feature of the basal keratinocyte cell death characteristic of the interface dermatitis of CLE.(11) IFN-I can also upregulate antigen presentation, co-stimulatory molecule expression, as well as promote the maturation of dendritic cells which may be important in the emergence of autoimmunity when engaged chronically.(12) IFN-I can also promote Th1 differentiation and cytotoxic skewing of T cells, the most abundant inflammatory cell type within CLE lesion, which may also be a fundamental aspect of CLE pathogenesis.(12,13)

The precise cause of the elevated IFN-Is of CLE remains an open question. Two well-known triggers of CLE, ultraviolet light and cigarette smoking, may cause cell injury or death which can result in the release of nucleic acids.(3–5) These nucleic acids may engage pattern recognition receptors resulting in the production of IFN-Is.(14) However, other sources of nucleic acids have been suggested as possible inducers, including noncoding RNAs and retroelements.(15,16) Retroelements are highly repetitive DNA sequences that comprise up to 50% of the human genome and though understudied, they have previously been implicated in autoimmune disorders as inducers of IFN-Is as well as potential autoantigens.(17)

In this study, using unbiased single-cell and spatial analyses of human CLE skin samples, we examined IFN-I and IFN signatures on the CLE microenvironment and probed potential sources of these signals.

## Results

We initially performed single cell RNA sequencing of skin biopsies from 3 healthy control (HC) donors and 5 patients with CLE, including non-lesional (NL) skin for a total of 12 samples and 35,079 cells (**Figure 1A; Supplementary Table 1; Supplementary** Figure 1A). Clustering with Uniform Manifold Approximation and Projection (UMAP) revealed 11 major cell types that were distinguished by selective expression of marker genes (**Figure 1B; Supplementary** Figure 1B). We first analyzed keratinocytes given their central role in the interface dermatitis of CLE (**Figure 1C**). Subclustering of keratinocytes showed 8 different subsets of keratinocytes including Basal, Suprabasal, Follicular, and Cycling cell populations (**Figure 1C; Supplementary** Figure 1C-D).

**Figure 1:**
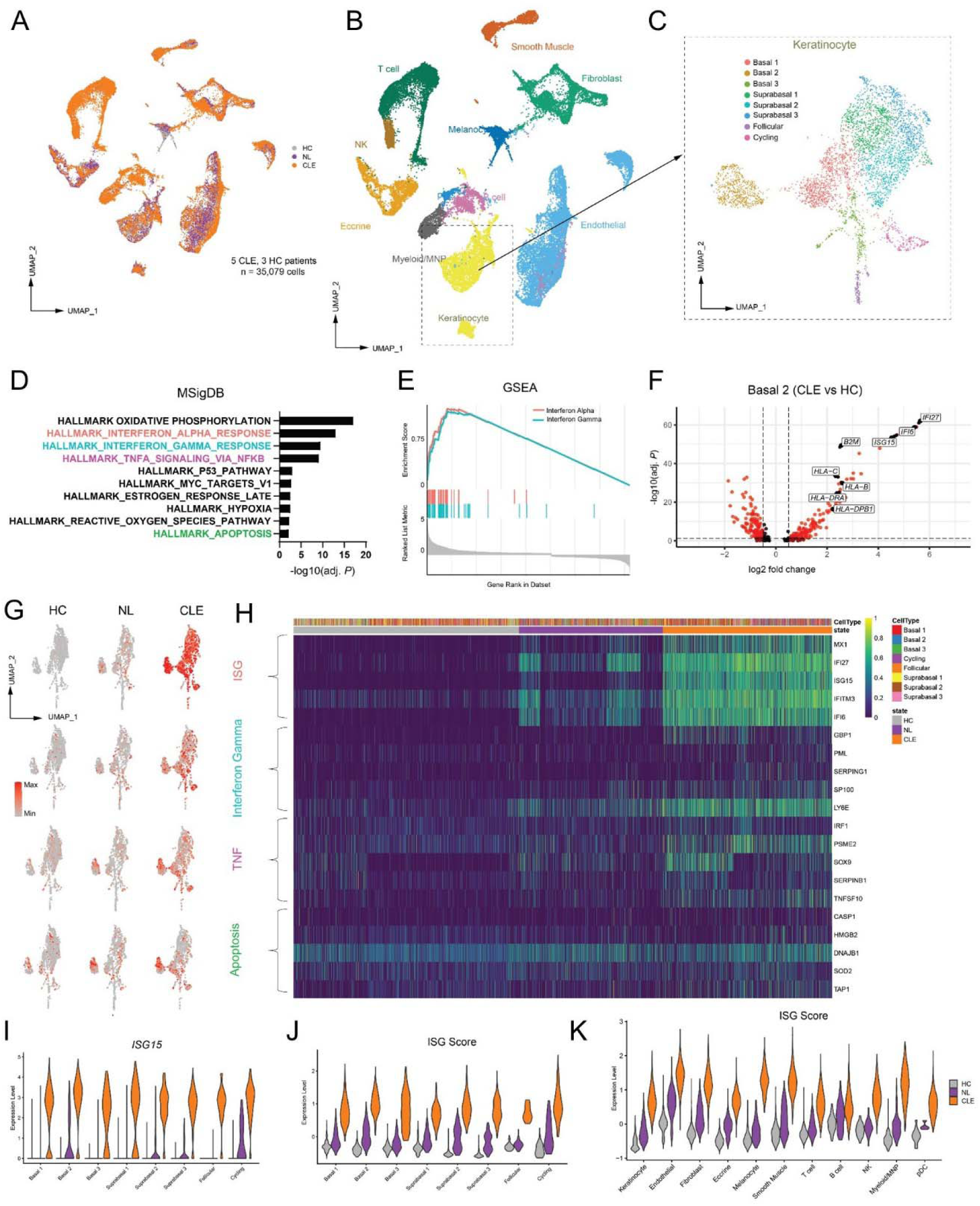
Single cell RNA sequencing of CLE reveals dominant type I interferon signature. A) UMAP plot of dataset of healthy control (HC), non-lesional (NL), and lesional (CLE) tissues. B) UMAP showing 11 distinct cell types in dataset. C) Subclustering of Keratinocytes with 8 cell types. D) MSigDB Hallmark Pathway analysis showing the top 10 upregulated pathways in lesional keratinocytes of CLE. E) Gene Set Enrichment Analyses (GSEA) showing enrichment of Interferon Alpha and Gamma pathways in lesional Keratinocytes. F) Volcano Plot of Basal 2 Keratinocyte clusters revealing upregulated interferon and antigen presentation target genes in lesional keratinocytes over controls. G) UMAP plots with overlayed gene signatures on Keratinocyte populations. H) Heatmap of Keratinocyte populations separated by cell type and state with 5 target genes from each signature represented (IFN-I, IFN-II, TNF, and Apoptosis). I, J) Violin plots showing upregulation of *ISG15* and IFN-I signature in CLE keratinocytes compared to NL and HC. K) Violin plot of IFN-I signature of all cell types broken down by cell state. UMAP, Uniform Manifold Approximation and Projection.

Unbiased MSigDB (18) Hallmark pathway analysis of CLE keratinocytes compared to HC revealed significant upregulation in oxidative phosphorylation and inflammatory pathways including response to interferon alpha, interferon gamma, TNF signaling, as well as apoptosis among the top 10 pathways (**Figure 1D**). The illustration of known pathways (Interferon Alpha) in CLE and those directly related to known histopathologic changes (apoptosis of keratinocytes) indicated our dataset contains core signals central to CLE biology. Gene Set Enrichment Analysis (19) (GSEA) confirmed the upregulation of interferon alpha and gamma pathways, with notable overlap of target genes but also a number of distinct targets, representing the simultaneous activation of both pathways in lesional keratinocytes (**Figure 1E**).

Our clustering analyses revealed the Basal 2 population of keratinocytes clustered separately from other keratinocytes, suggesting distinct biology (**Figure 1C**). When comparing gene expression of CLE and HC in Basal 2 Keratinocytes, we observed these cells to markedly upregulate antigen presentation machinery, including MHC I and MHC II genes (**Figure 1F; Supplementary** Figure 1E**-F**). Expression of MHC II genes was previously seen in the keratinocytes of healing wounds (20) and other inflammatory contexts;(21) upregulation of MHC II has been observed in differentiated keratinocytes of CLE(20) but not basal keratinocytes. Evaluation of IFN-I, IFN-II, TNF, and Apoptosis gene signatures (see methods) in HC, NL, and CLE indicated some degree of elevated signals in NL skin, especially ISGs (**Figure 1G-H**), which has been observed in prior work.(22) The elevation of *ISG15* as well as the IFN-I signature in CLE was pervasive across all keratinocyte clusters, with a more modest elevation of NL compared with HC skin. Yet, we did not observe IFN-I signatures in any NL keratinocyte clusters that were higher than that seen in the lesional CLE, as reported recently (**Figure 1I**-**J**).(22) A similar pattern of high IFN-I signature in CLE and modest to moderate elevated IFN-I signature in NL samples was seen across all cell types (**Figure 1K**).

To garner more insights into how these 11 different cell types may communicate in different tissue states (HC, NL, and CLE), we performed ligand-receptor analysis using CellChat.(23) Throughout all three states, fibroblasts had the highest outgoing signal strength, indicating the highest potential for recruitment of cells into the skin (**Figure 2A**-**C**). A combination of soluble and structural signals ranked in the top 15 of interactions in CLE, with MIF signaling ranked the highest across all cell types (**Figure 2C**). While Myeloid/Mononuclear Phagocyte (MNP) and NK cells had among the highest incoming interaction strengths across all three states, pDCs were uniquely increased in the CLE state compared to NL and HC tissue. Historically, pDCs were presumed to be the major producers of IFN-I in CLE. However, recent studies have shown that pDCs display an exhausted phenotype in lupus skin, suggesting their accumulation may not directly correlate with functional type I IFN production.(7)

**Figure 2:**
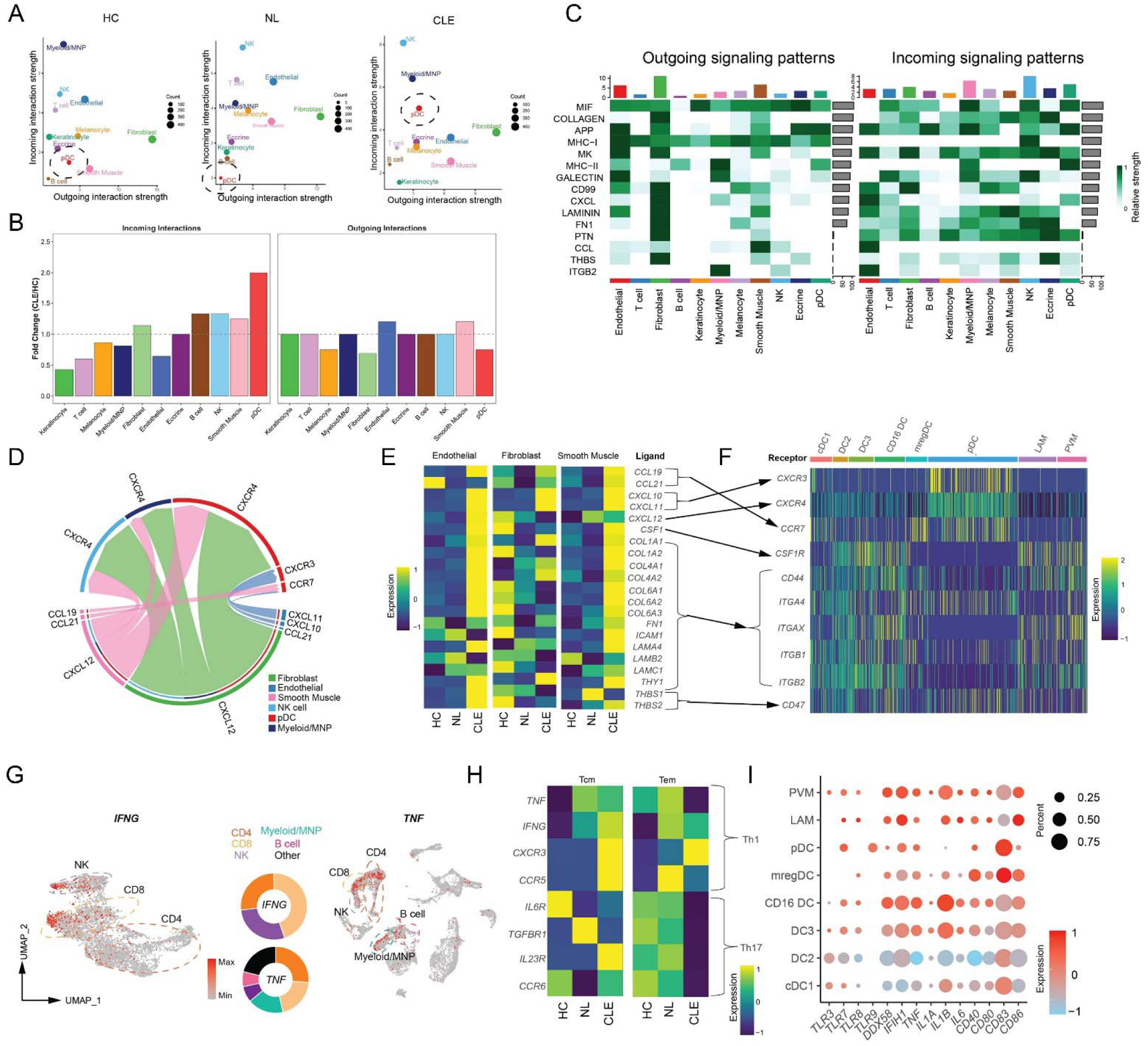
Cell interaction analysis of stromal and inflammatory cells in CLE. A) Ligand-Receptor analysis with CellChat demonstrating cell type interactions by each tissue state. Black dashed circles identify pDCs. B) Ratio of interaction strengths between HC and CLE for different cell types. C) Heatmap of cell type interactions with the top 15 pathways included. D) Chord diagram of CCL and CXCL pathways annotated by the top 3 “sender” and “receiving” cell types. E) Heatmap of ligands identified by ligand-receptor analysis separated by cell type among the top 3 “sender” cell types. F) Heatmap of MNP populations from CLE tissues showing expression of upregulated ligand receptors by different cell types. G) UMAP plots highlighting cells with detectable *IFNG* and *TNF* expression; donut plots show the percentage of cell types. H) Heatmap of CD4 T effector memory and central memory populations with expression of Th1 and Th17 markers, separated by tissue state. I) Bubble plot of MNP populations from CLE tissues with expression of pattern recognition receptors, inflammatory markers, and activation markers. Color indicates relative expression levels between cell types in CLE and the size indicates the percentage of cells per cluster expressing the particular gene.

Given the known roles of inflammatory chemokines in mobilizing immune cells into inflamed tissues, we further examined the interactions between the top 3 “sender” and “receiver” cell types in CLE tissues among these pathways (**Figure 2D**). This analysis indicated the importance of the *CXCR3*, *CXCR4*, and *CCR7* chemokine axes in the recruitment of pDCs into CLE skin, whereas *CXCR4* alone was implicated in the accumulation of Myeloid/MNP and NK cells. We next evaluated the expression of specific ligands suggested from this analysis across cell states and compared this with the expression of their receptors on Myeloid/MNP and pDC cells (**Figure 2E**). For many of the studied ligands, we observed their specific upregulation in the CLE state, especially in endothelial and smooth muscle cell types. In CLE tissues, *CXCR3* and *CXCR4* receptor expression were most abundant in pDCs, and *CCR7* was most abundant in both pDCs and mregDC populations, the latter suggestive of their maturation status (**Figure 2F**). We further evaluated *CXCL10* and *CXCL11* expression in all cell types and found it uniquely upregulated in CLE tissues and most abundant in endothelial and Myeloid/MNP cells (**Supplementary** Figure 2A-B), whereas *CXCL12* expression was largely stable across tissue states and most abundant in endothelial, fibroblast, and smooth muscle cells (**Supplementary** Figure 2C).

We further probed for the sources of IFN-γ and TNF. The majority of lymphoid cells producing *IFNG* were CD8 T cells (45%) followed by NK cells (28%), with the remainder produced from CD4 T cells (27%, **Figure 2G**). The majority of cells producing *TNF* were also T cells (46%), followed by Myeloid/MNP (17%), NK cells (8%), and B cells (7%). These studies indicate the relative importance of CD4 and CD8 T cells producing key cytokines in the CLE microenvironment, influencing CLE keratinocyte biology. In addition to type I interferons lowering the threshold for apoptosis in keratinocytes,(11) IFN-γ has also been shown to promote apoptosis in keratinocytes of lichen planus,(24) another dermatosis featuring interface dermatitis like CLE. Moreover, the interaction of IFN-γ and TNF has been shown to promote synergistic cell death in keratinocytes and other cell types.(25,26) Thus, our data implicate T cell elaboration of key cytokines and cell death in the interface dermatitis of CLE.

Prior studies have demonstrated evidence of Th1 and/or Th17 skewing present in SLE and CLE.(3,27) We first subclustered lymphoid cells into 9 discrete clusters (**Supplementary** Figure 2D-**E**). Examination of CD4 central memory (Tcm) and CD4 effector memory T cell (Tem) populations for markers of Th1 and Th17 cells showed clear skewing for Th1 cells in Tcm and better evidence for a Th1 compared to Th17 phenotype in Tem (**Figure 2H**). *IL17A, IL17E, and IL17F* were detected in very few cells in our dataset. Analyses of CD8 T cells and NK cells revealed elevated IFN-I signatures as well as increased activation in CLE (**Supplementary** Figure 2F-I).

We further examined MNP populations in CLE. After subclustering MNP populations (**Supplementary** Figures 2J-K**, 3-5**), we evaluated their expression of pattern recognition receptors (PRRs), key inflammatory cytokines, and markers of activation (**Figure 2I**). pDCs expressed the highest levels of *TLR7* and were the only cell type to express *TLR9*, but expressed neither *TLR3* nor *TLR8*, as expected. RIG-I (encoded by *DDX58*) and MDA5 (encoded by *IFIH1*) were expressed by all MNPs, with the highest amounts in perivascular macrophages (PVM), CD16+ DCs, and DC3s. Similarly, these three cell types expressed the highest levels of inflammatory mediators *TNF*, *IL1A*, *IL1B*, and *IL6*. Among *CD40*, *CD80*, *CD83*, and *CD86* – all costimulatory molecules and markers of MNP activation – CD83 was expressed at the highest levels in mregDCs and pDCs. Thus, these data indicate the inflammatory microenvironment of CLE skin includes a variety of MNP cell types expressing a complement of PRRs poised for sensing innate signals, including highly mature pDC and mregDC populations expressing elevated levels of *CCR7* and *CD83*. Additionally, these results further highlight a role for monocyte-derived MNPs (28,29) in CLE as recently described,(22) with PVM, CD16 DCs, and DC3s demonstrating the most inflammatory phenotype among MNP populations.

Given that our cell interaction analysis suggested an important role of fibroblasts signaling to immune cells in the skin of CLE, we performed more detailed studies of our fibroblast populations. Sub-clustering analysis revealed a total of 12 distinct clusters (**Figure 3A**-**B**). We observed an expansion of CLE fibroblasts in F1, F4, and F7 clusters (**Figure 3B**). *GREM1* and *PLA2G2A* marked the F4 and F7 (as well as F1 and F6) expanded clusters, respectively (**Figure 3C**). Given their UMAP proximity and shared marker gene expression, our subsequent analyses of *PLA2G2A*+ fibroblasts included the F1, F6, and F7 clusters. Pseudotime trajectory analysis of fibroblast clusters (30,31) revealed a developmental path ending with these unique fibroblast clusters F4, F7, and F6 (**Figure 3D**-**E**). With the unique expansion of these clusters and their association with elevated interferon signaling as seen by elevated *ISG15* expression (**Figure 3D**), these data suggest the inflammatory milieu of CLE promotes an altered differentiation trajectory leading to the accumulation of *GREM1*+ and *PLA2G2A*+ fibroblasts.

**Figure 3:**
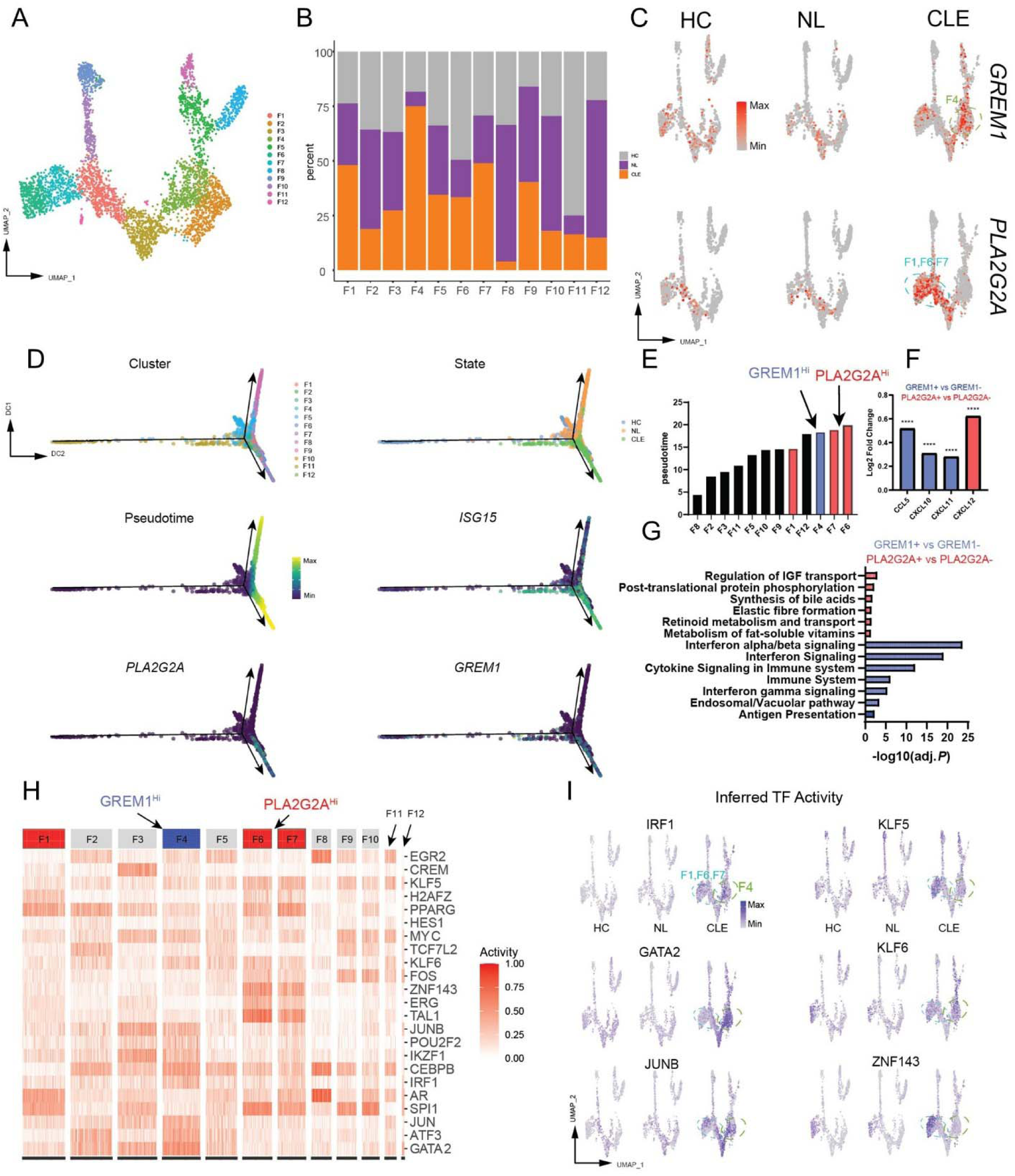
Inflammatory CLE environment reprograms the fibroblast lineage. A) UMAP plot of detailed fibroblast subclustering. B) Stacked bar plot of percentage breakdown of tissue state in different fibroblast groups. C) UMAP plots of *GREM1* and *PLA2G2A* expression separated by cell state. D) Diffusion maps of cells labeled by cluster identify, tissue state, pseudotime, as well as *ISG15*, *GREM1*, and *PLA2G2A* expression. E) Cell clusters rank ordered by median pseudotime. F) Bar plot of chemokines upregulated in labeled clusters, **** *P* < .0001, Wilcoxon Rank Sum test. G) Reactome pathway analysis showing upregulated pathways in *PLA2G2A*+ and *GREM1*+ fibroblasts. H) BITFAM analysis of fibroblast clusters showing different patterns of transcription factor activity. I) UMAP plots of inferred transcription factor

We next compared the expression profiles of *GREM1*+ vs. *GREM1*- and *PLA2G2A*+ vs. *PLA2G2A*-fibroblasts in CLE skin. *GREM1*+ fibroblasts had increased expression of *CCL5*, *CXCL10*, and *CXCL11*, whereas *PLA2G2A*+ fibroblasts had increased expression of *CXCL12* (**Figure 3F**). We performed Reactome (32,33) pathway analysis of differentially expressed upregulated genes in these *GREM1*+ and *PLA2G2A*+ populations for more insights into their potential function. This showed an elevated IFN-I signature in *GREM1+* fibroblasts and evidence of increased extracellular matrix and collagen synthesis in *PLA2G2A+* fibroblasts (**Figure 3G**). For additional study of the transcriptional signatures of these unique fibroblasts, we performed Bayesian inference transcription factor activity model (BITFAM) analysis (**Figure 3H**).(34) Examination of individual transcription factors like IRF1, GATA2, and JUNB revealed elevated activity in either or both *PLA2G2A*+ and *GREM1*+ fibroblast clusters compared to NL and HC fibroblasts (**Figure 3I**).

Recent studies have suggested that blockade of type I interferon signaling with Anifrolumab may be a highly effective therapy for CLE.(10,36) For further insight into how type I interferon blockade may result in improvement in CLE disease, we performed 10x Visium spatial sequencing of a patient with subacute cutaneous lupus (SCLE) before and after treatment with Anifrolumab in addition to control and discoid lupus (DLE) samples. Eight weeks after treatment with Anifrolumab, the patient’s skin disease was largely remitted without active inflammation present (**Figure 4A**). Pre- and post-treatment biopsies were site-matched and directly adjacent to the forearm. Clustering analysis and UMAP plots revealed 12 clusters labeled by their predominant cell type within the ∼50 µm cell spots (**Figure 4B**). Pre-Anifrolumab treatment, the CLE sample had 4 unique clusters that were not present in post-treatment CLE and healthy controls (**Figure 4C**). This included a cluster of inflammatory keratinocytes (KC_Inflam), a cluster of mixed cell types including myeloid cells (Mixed_Myeloid), as well as two clusters of fibroblasts – *GREM1+* and *PLA2G2A+* fibroblasts that were previously identified in our single cell dataset. This suggests that the accumulation of these unique fibroblasts is dependent on type I interferon signaling, as indicated in our prior analysis.

**Figure 4:**
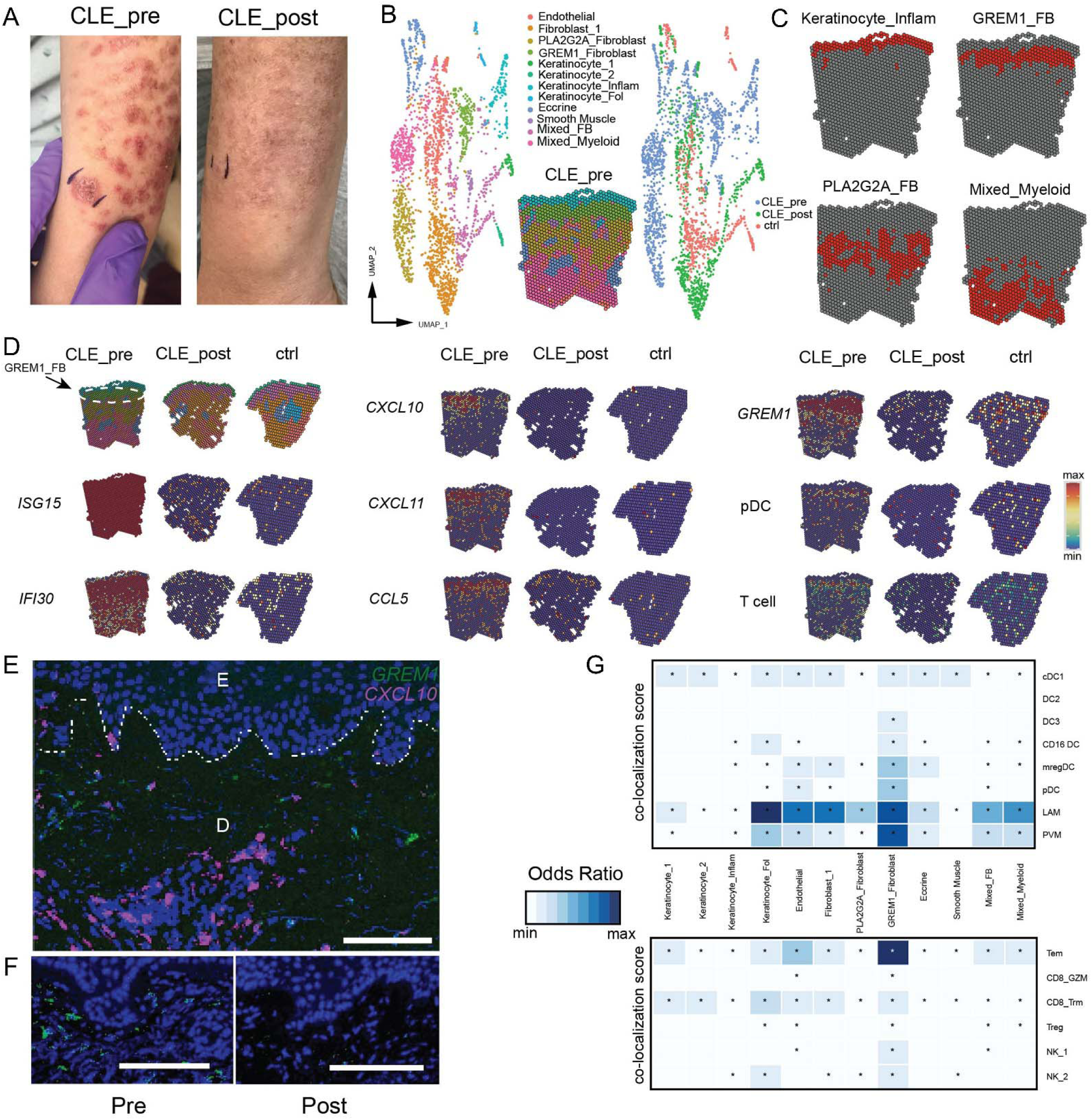
Spatial sequencing suggests pathogenic importance of type I interferon-dependent GREM1 Fibroblasts. A) Pre- and post-treatment photos (8 week interval after Anifrolumab) of patient with subacute cutaneous lupus showing near complete resolution in disease activity. B) UMAP clustering of 10x Visium spatial sequencing of skin biopsy tissues from pre-treatment active disease (CLE_pre), post-treatment (CLE_post), and healthy control (ctrl) tissues. C) Unique clusters present only in CLE active disease sample. D) Spatial plots of gene expression for the labeled genes or gene signatures for a particular cell type (pDC, T cell). E) In situ hybridization of *GREM1* (green) and *CXCL10* (magenta) in CLE skin. E labels pre-treatment epidermis and D for dermis, with the dotted line indicating the demarcation. Magnification 200X, scale bar = 100 µM. F) Pre- and post-Anifrolumab (12 week) site-matched biopsy displaying the dermal-epidermal junction of discoid lupus stained with *GREM1*. Magnification 200X, scale bar = 100 µM G) Co-localization score for localizing cell type specific signatures in spatial sequencing data. Upper panel shows myeloid signatures from our single cell dataset (labels on right) and lower shows lymphoid signatures. Quantified using Multimodal intersection analysis (see methods). * indicates statistically significant enrichment (*P* < 0.05) and color intensity maps to elevated odds ratios.

Visual inspection of the localization of spatial clusters revealed a surprising finding – the *GREM1*+ fibroblast (GREM1_FB) population was localized to the superficial dermis, adjacent to inflammatory keratinocytes and directly at the site of pathology in CLE (**Figure 4B**). Analysis of *ISG15* and *IFI30* revealed diffuse activity across the entire tissue sample that was only present at low levels in post-treatment and control tissue (**Figure 4D**). Evaluation of the chemokines *CXCL10*, *CXCL11*, and *CCL5* demonstrated robust elevation in the papillary and superficial reticular dermis of pre-treatment tissue compared to post-treatment and control skin tissue. In accordance with the presence of GREM1+ fibroblasts only in pre-treatment tissue, *GREM1* expression was markedly decreased in post-treatment and control tissues. pDC (*IL3RA*, *LILRA4*, *CLEC4C*) and T cell (*CD3E*, *CD3D*, *IL7R*) signatures derived from our single cell analysis revealed co-localization of pDCs and T cells in the papillary dermis with *GREM1+* fibroblasts at the site of tissue pathology in CLE. For confirmation of *GREM1*+ fibroblasts and their localization, we performed in-situ hybridization which revealed *GREM1*+ cells in the superficial dermis of CLE tissues, some of which expressed *CXCL10* (**Figure 4E**). We also show that *GREM1*+ fibroblasts were present in both subacute cutaneous lupus (**Figure 4D**) and discoid lupus (**Supplementary** Figure 6A) with the same localization, associated chemokines (*CXCL10*, *CCL5*), and associated cell types (pDC, T cells). *GREM1*+ cells were also confirmed to be present in the superficial dermis of an additional patient with DLE prior to Anifrolumab and absent in a site-matched biopsy 12 weeks post-initiation (**Figure 4F**).

For further resolution on inflammatory cell localization within our CLE spatial transcriptomics dataset, we performed multimodal intersection analysis (MIA) which enables deconvolution of the cell types present in the ∼50 µm spots of potentially overlapping cell types (**Figure 4G**, 36). Through this analysis, MIA informs which cell types co-localize. Analysis of 8 different MNP cell types with signatures derived from our single cell analysis showed the most enrichment in *GREM1*+ fibroblasts; every MNP signature aside from DC2 was enriched in *GREM1*+ fibroblasts (**Figure 4G**). Similarly, analysis of 6 lymphoid signatures demonstrated the most enrichment in *GREM1*+ fibroblasts. These analyses further implicate *GREM1*+ fibroblasts as an important hub for the initiation or maintenance of inflammation in CLE.

Mucin deposition is a histopathologic hallmark of CLE. Given that PLA2G2A+ fibroblasts are observed in both CLE and scleroderma(35), another autoimmune skin disease with mucin deposition, we examined mucin-associated gene expression across fibroblast populations. Analysis of individual mucin-related genes (*GALNT1, GALNT2, MUC1, MUC3A, MUC6, TFF3*) revealed elevated expression in all fibroblast subtypes in CLE compared to other tissue states (**Supplementary** Figure 7). Notably, GREM1+ fibroblasts showed the highest aggregate mucin gene signature score among fibroblast populations, suggesting these cells may be the major contributor to the characteristic mucin deposition seen in CLE (**Supplementary** Figure 8).

Given the pervasive elevation of type I interferons in CLE, we investigated potential associations with retroelement expression. Prior work from our group and others has implicated the deregulated expression of retroelements in the pathogenesis of SLE;(38–42) elevated retroelement expression has the potential to engage nucleic acid pathogen recognition receptors to stimulate type I interferon expression. Retroelements are repetitive DNA sequences that include endogenous retroviruses (ERVs) as well as Long Interspersed Nuclear Elements (LINEs) and Short Interspersed Nuclear Elements (SINEs). Whereas only 1-2% of the human genome encodes protein-coding genes, these retroelements comprise roughly 50% of our genome yet their role as a potential trigger of autoimmunity remains relatively unexplored; in homeostatic conditions, expression of retroelements is typically epigenetically silenced. To further examine retroelements, we used scTE(43), a computational pipeline specifically designed to identify and quantify transposable element expression from single-cell RNA sequencing data, to map retroelements in our single-cell dataset.

scTE analysis revealed deregulated expression of retroelements across multiple cell types in both NL and CLE tissues (**Figure 5A**). Though many retroelements were only differentially expressed in certain cell types, some were elevated across multiple cell types including L2b (LINE2 element), L1PA7 (LINE1 element), AluYb8, and AluYb9 (SINEs, **Figure 5A, B**). A more detailed examination of endothelial cells and fibroblasts revealed that the overwhelming majority of retroelements were upregulated (**Figure 5B**). LINEs and SINEs were the predominant upregulated classes of retroelements, with the remainder being ERVs and DNA transposons. We additionally evaluated the known epigenetic regulators of retroelements across the same cell types. We observed cell type-specific expression of members of the TRIM28, NuRD, and HUSH complexes that were frequently deregulated in NL and CLE tissues compared to HC. Notably, TRIM28 and SETDB1, two critical regulators of retroelement silencing,(44–49) were downregulated across most cell types in CLE (**Supplementary** Figure 6B).

**Figure 5:**
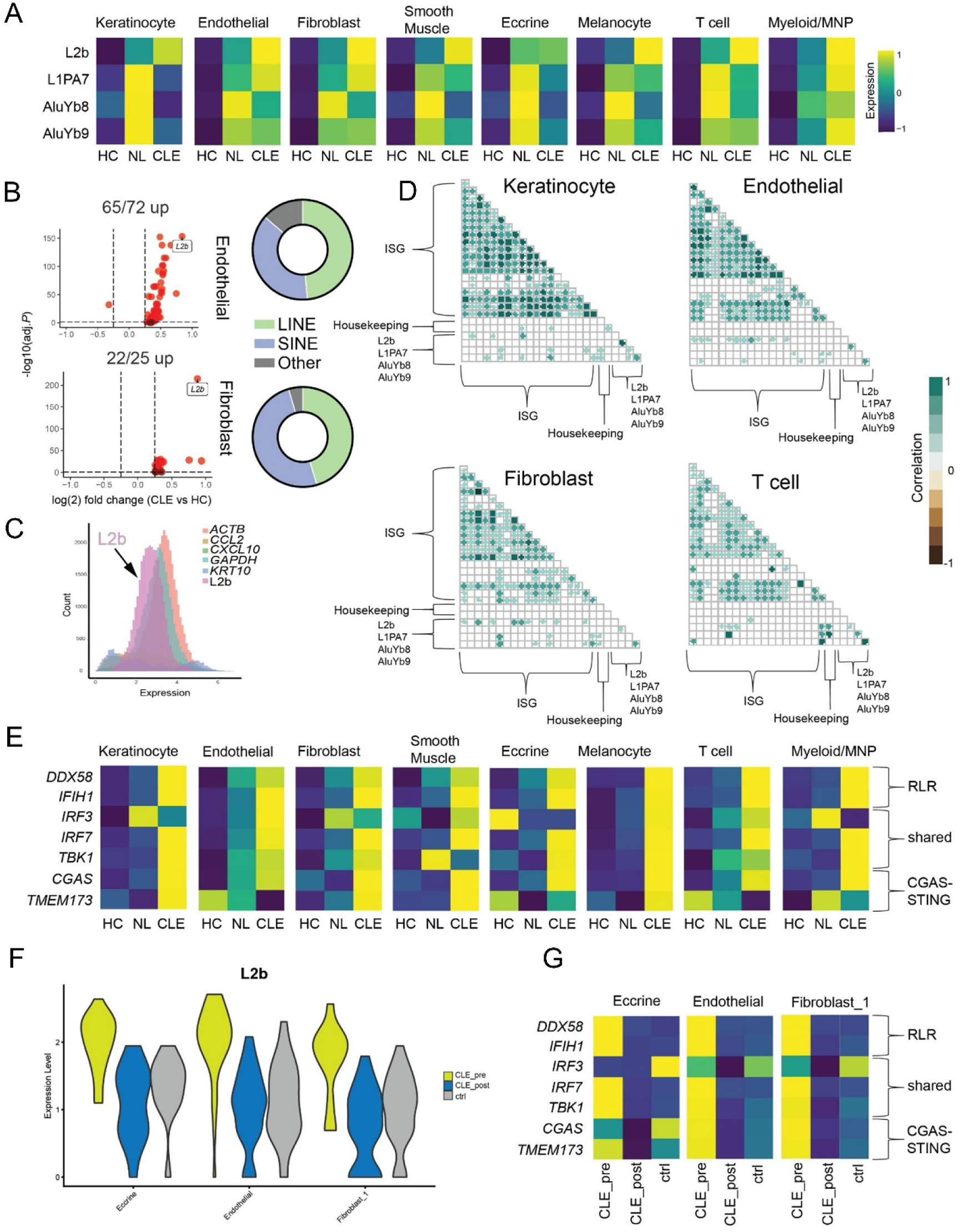
Deregulated expression of transposable elements correlates with interferon signature in CLE. A) Heatmap of scTE analysis of skin dataset reveals elevated retroelement expression across multiple cell types in NL and CLE skin. B) Volcano plot of Endothelial and Fibroblast clusters (CLE vs HC). Donut plots show distribution of statistically significant upregulated retroelements (adjusted *P* < 0.05). C) Histogram of cell counts (y axis) and expression levels (x axis) of selected genes. D) Correlation matrix showing statistically significant correlations (*P* < .0001) between ISG expression and retroelements among labeled cell types. Non-significant correlations remain blank. E) Heatmap of RLR and cGAS-STING pathway gene expression among different cell types and tissue states in CLE. F) Violin plot demonstrating decrease in L2b retroelement levels after treatment with CLE treatment Anifrolumab, comparable to healthy control levels. These are fibroblast clusters identified in Figure 4B. G) Heatmap of RLR and cGAS-STING pathway gene expression pre and post-Anifrolumab in CLE.

In multiple cell types, L2b was the most abundant differentially expressed retroelement in CLE (**Figure 5A-B**). We compared the abundance of L2b to other more abundant housekeeping (*ACTB*, *GAPDH*) and cell type-specific (*CXCL10*, *CCL2*, *KRT10*) genes, observing that L2b expression levels were similar to that of the highly expressed housekeeping genes *ACTB* and *GAPDH* (**Figure 5C, Supplementary** Figure 9). L2b levels were upregulated in all CLE cell types relative to HC (**Supplementary** Figure 10). Given that retroelements are well-described inducers of the type I interferon response,(17) we evaluated the correlation of differentially expressed retroelements with the ISG response across multiple cell types. Using housekeeping genes (*ACTB*, *GAPDH*) as the “background” level of correlation between ISGs and the retroelements L2b, L1PA7, AluYb8, and AluYb9, we observed elevated, statistically significant associations in at least 3 of the 4 retroelement metagenes with multiple ISGs in keratinocytes, endothelial cells, fibroblasts, and T cells (**Figure 5D**). Though most correlations were mild to moderate (0.2 – 0.6), in T cells some of the correlations were higher (0.8 – 1.0). These data indicate that retroelements, which may be abundant in the skin and upregulated in CLE, could be a potential driver of the elevated type I interferon responses seen in these tissues.

The RIG-I like receptors (RLR) and cGAS are innate immune sensors that are broadly expressed and critical components of the antiviral response by binding viral RNAs (RLRs) or cytosolic DNA (cGAS), respectively, to stimulate a type I interferon responses.(50,51) Retroelements have been shown to promote type I interferon responses through RLRs as well as cGAS-STING.(17,52) We examined the expression of RLR and cGAS-STING pathway genes across cell types and tissue states, revealing marked upregulation across many cell types in CLE tissues (**Figure 5E, Supplementary** Figure 11). Overexpression of RLRs and cGAS can increase a cell’s innate responses to nucleic acid ligands,(53,54) and these sensor genes themselves are enhanced by type I interferons, suggesting that CLE tissues may be primed for heightened inflammatory responses to nucleic acid stimuli and permissive of a positive feedback loop to maintain high ISG state.

Given this question of feedback loops with a possible role in the persistent inflammation driving CLE lesions, we next asked whether Anifrolumab treatment alters retroelement and innate immune receptor expression. We examined cell types abundant in pre- and post-treatment CLE as well as control tissue. Notably, L2b, the most abundant differentially expressed retroelement present in our CLE analysis, decreased to control levels with Anifrolumab treatment (**Figure 5F**). In addition, Anifrolumab normalized the expression of RLR and cGAS-STING pathway genes, as expected (**Figure 5G**). These data suggest that 1) type I interferon signaling is required to maintain the expression of certain retroelements and 2) Anifrolumab’s clinical efficacy may, in part, be due to breaking this positive feedback loop of elevated sensor gene expression responding to elevated expression of nucleic acid ligands.

## DISCUSSION

In this study, we examined lesional and non-lesional biopsies of patients with CLE using single cell RNA and spatial sequencing approaches. We show a dominant IFN-I signature that is pervasive across all cell types with a milder elevation in non-lesional skin. Pathway analysis also revealed robust activation of TNF and IFN-γ pathways in CLE keratinocytes, indicating the elaboration of these key cytokines at the interface dermatitis of CLE. We delineate the most abundant sources of these cytokines in lesional skin as CD4 and CD8 T cells for *TNF* and CD8 and NK cells for *IFNG*, further establishing the role of lymphocytes in the immunopathology of CLE. Cell communication analysis revealed important interactions between pDCs and stromal cell populations utilizing the CXCR3, CXCR4, and CCR7 chemokine axes, suggesting these pathways may be valuable targets to evaluate for treatment of CLE in the future.

There are several important limitations to our study. Our high-dimensional single-cell and spatial analyses utilize a relatively small cohort of patient samples. Additionally, our pre- and post-biopsy analyses are based on one patient and may represent indirect findings of resolved skin inflammation rather than direct effects of type I interferon in the skin. Our retroelement analysis provides family level expression without locus specific details, and the correlations observed between retroelements and interferon signatures may reflect either cause or consequence of interferon activation. The most dysregulated pathway we observed was oxidative phosphorylation, suggesting mitochondrial stress could be an important upstream driver of interferon responses. However, our study nonetheless provides several new insights into CLE pathogenesis including the reprogramming of local fibroblasts to an inflammatory phenotype, a central role of local T cells in elaboration of pathology, and type I IFN correlation with retroelement expression.

Given our cell interaction analysis indicated particular importance of CLE fibroblasts as a source of ligands, we performed a more detailed study of fibroblasts in CLE. We observed that the IFN-I-rich microenvironment of CLE was associated with two unique populations of fibroblasts, *GREM1*+ and *PLA2G2A*+ fibroblasts. *GREM1*+ fibroblasts expressed higher levels of *CXCL10*, *CXCL11*, and *CCL5* whereas *PLA2G2A*+ fibroblasts expressed higher levels of *CXCL12* – all likely important ligands in sustaining the leukocyte infiltration into the CLE skin. With recent studies indicating the therapeutic efficacy of targeting pDCs in CLE,(55) these data implicate the *CXCR3* and *CXCR4* pathways in the recruitment of pDCs into CLE skin and by extension potential therapeutic targets.

By utilizing patient samples pre- and post-treatment with the IFNAR-blocking antibody Anifrolumab, we show these unique fibroblast populations are specific to CLE and dependent on type-I interferon mediated inflammation. Our spatial analyses reveal a critical localization of *GREM1*+ fibroblasts to the papillary dermis adjacent to the inflammatory keratinocytes in CLE, suggesting they may have a pivotal role in immunopathology. *GREM1*+ fibroblasts have been observed in skin cancers as well as fibrotic lung diseases.(56,57) GREM1 is a BMP antagonist that, in a number of contexts, promotes mesenchymal fibroblast phenotypes.(58) Notably, recent studies have uncovered a role of *GREM1*+ fibroblast cells in secondary lymphoid organs, where they are critical for dendritic cell localization and homeostatic function.(59) Though more studies are necessary, these data support a role for GREM1+ fibroblasts in the recruitment or retention of inflammatory cell populations in the papillary dermis of CLE, intimately associating them with the tissue damage observed at the dermal-epidermal junction. In addition, we also observed *PLA2G2A* fibroblasts deeper in the dermis of patients that notably with predicted functions of collagen synthesis and extracellular matrix functions, suggesting a role in fibrosis.

PLA2G2A fibroblasts are also present in scleroderma.(35) It is possible these are the fibroblasts chiefly responsible for the fibrotic scarring observed in late-stage DLE lesions. The presence of PLA2G2A+ fibroblasts in both CLE and scleroderma points to shared stromal alterations in autoimmune skin diseases. Our analysis revealed broadly elevated mucin-associated gene expression across fibroblast populations in active disease, with GREM1+ fibroblasts showing particularly high expression of this gene program. This finding provides a potential mechanistic link between specialized fibroblast populations and the mucin deposition characteristic of CLE and other autoimmune skin conditions. Understanding how these distinct fibroblast populations contribute to shared disease features may reveal common pathways that could be therapeutically targeted.

While our study leverages both single-cell and spatial transcriptomics to understand CLE pathology, additional computational analyses integrating these datasets could yield further insights. For example, examining the gene regulatory networks active in spatially-defined GREM1+ fibroblast populations could help identify key transcriptional programs driving their inflammatory phenotype. Similarly, systematic comparison of ligand-receptor pairs identified in our single-cell analysis with their spatial distribution could better define the molecular mediators of immune cell recruitment to these inflammatory niches. These integrated analyses across different disease subtypes and treatment timepoints could provide a more comprehensive understanding of how cellular networks in the skin adapt during disease progression and resolution.

Finally, we examined retroelement expression patterns in our dataset. While retroelements have been associated with autoimmune disorders and can potentially drive type I interferon responses, their precise role remains unclear. (17,40,41) We observed correlations between retroelement expression and interferon signatures across multiple cell types. The tissue microenvironment of CLE showed elevated expression of nucleic acid sensing pathways including RLRs and cGAS-STING, which could potentially respond to various triggers including retroelements but also other sources such as mitochondrial DNA released during cellular stress. In addition, our findings indicate that blocking type I interferon signaling with Anifrolumab reduces both retroelement and innate immune sensor expression, suggesting these pathways may participate in a feedback loop that maintains inflammation in CLE. Though additional studies are needed to fully elucidate these pathways, our findings point to potential therapeutic strategies in CLE that could target not only type I interferon signaling but also the molecular and cellular networks that maintain inflammatory feedback loops in the skin.

## Author Contributions

J.R.G. and A.I. were responsible for designing the study, analyzing data, and writing the manuscript. J.R.G and E.R.B conducted the experiments. Y.K., and M.V. assisted in data analysis and manuscript preparation. C.J.K performed the histopathologic analysis and tissue procurement. B.K. and W.D. assisted with patient recruitment and IRB preparation. A.J.L, S.R., and F.K. contributed to patient recruitment.

## Acknowledgments

This work was supported in part by the Lupus Research Alliance Distinguished Innovator Award and the Howard Hughes Medical Institute. J.R.G. received research and salary support from the Yale Physician Scientist Development Program and the Dermatology Foundation for career development awards. We thank Keck Microarray Shared Resource and Yale Center for Genome Analysis at Yale University for their assistance with 10x Genomics single cell RNA-sequencing services.

## Methods

### Sex as a Biological Variable

Nearly every enrolled patient in our study was of female gender given the significant sex bias in cutaneous lupus. These results are still generalizable given the prevalence of CLE in women, though the applicability to men with cutaneous lupus is unknown.

### Skin digestion for single cell RNA sequencing

4 or 5 mm punch biopsies were taken skin and digested as follows. Biposies were bisected into four pieces. Dispase II (Sigma) was added at a concentration of 10 mg/mL in RPMI supplemented with 10% Fetal Bovine Serum, 1% L-Glutamine (Gibco), and 1% Penicillin/Streptomycin (Gibco). After 45 minutes of incubation in 1 mL of digestion media, the tissue was placed on a 6mm dish and cut into finer pieces with a razor. The media and tissue were placed in 5-10 mL of 0.5 mL/mL Liberase TH (Sigma) with 40 ug/mL DNase I (Roche) and digested another 45 minutes. Tissue and media were then moved through a 70 uM strainer (Fischer Scientific) and counted using a Countess III (ThermoFisher). Cells were then suspended in 0.04% Bovine Serum Albumin (BSA) and phosphate-buffered saline (PBS) and taken to the Yale Center for Genome Analysis for 10x Genomics single cell RNA library preparation and sequencing.

### Single cell RNA sequencing and analysis

Yale Center for Genome analysis (YCGA) assisted with 10x Genomics single cell sample preparation, including library construction and sequencing. Sequencing libraries were constructed using the 10x Genomics 5’v2 kit. Samples were sequenced at a minimum depth of 20,000 read pairs per cell. Cell Ranger analysis pipelines were then utilized for alignment, gene counts, and filtering.

Analysis of single cell RNA sequencing data was performed using the Seurat(60) package (version 4.3.0) available with the R statistical programming environment (version 4.2.3). For quality control, cells with less than 200 or over 10,000 unique genes as well as cells that have greater than 5% mitochondrial reads were removed. Samples were then log-normalized and integrated using Seurat.(61) A subset of 2,000 highly variable genes by cell-to-cell variation were selected for downstream clustering and visualization analysis. Cells were clustered using a K-nearest neighbor graph-based clustering approach incorporated in the Seurat package. Non-linear dimension reduction method UMAP (Uniform Manifold Approximation and Projection) was used for visualizing the cells and clusters in 2D space. CellChat was used for cell-cell communication inferencing.(62) BITFAM was used for transcription factor activity analysis.(34) Gene set enrichment and pathway analyses were performed using ClusterProfiler(63) v4.6.2 and the GProfiler(33) web portal. Visualization plots and graphs were generated using Seurat and DittoSeq v1.14.0.

### Single cell RNA gene signatures/scores

For gene signatures/scores, the AddModuleScore function in Seurat was used. The IFN-I/ISG score was generated from a list of 20 interferon stimulated genes (*MX1, BST2, IFITM1, IFITM2, IFITM3, IFI44L, IFI6, IFI27, LY6E, IFIT1, IFIT2, IFIT3, ISG20, ISG15, CXCL9, CXCL10, CXCL11, OAS1, OAS2, OAS3*). This was also the gene list used in **Figure 5** for correlation analysis. TNF(64), Interferon Gamma(65), and Apoptosis(18) signatures were derived from prior publications. CD8 T cell and Natural Killer cell activation signatures were also generated based on prior work.(66,67)

### Multimodal Intersection Analysis

Multimodal intersection analysis was performed similar to that done in prior publications.(37) Cell type signatures from single cell analysis were generated by taking the top 20 statistically significant marker genes (as determined using the default parameters of the FindMarkers function in Seurat) from labeled cell types (see **Figure 1**) and calculating enrichment in the spatial cluster gene sets (see **Figure 4**) using GeneOverlap v1.34.0.

### scTE analysis

For retroelement analysis, scTE(43) version 1.0 was used. The hg38 genome index was built by scTE command “scTE_build -g hg38”. BAM files generated from the 10x Genomics CellRanger program were used as input for scTE. The BAM files were pre-filtered to retain only the mapped reads with proper CB tags (cellular barcode sequence that is error-corrected and confirmed against a list of known barcode sequences) and proper UB tags (molecular barcode sequence that is error-corrected among other molecular barcodes with the same cellular barcode and gene alignment). This prefiltering was done with Samtools (version 1.16).(68) This output was then loaded into Seurat and additional filtering was performed to only include cells that met our quality control criteria described above.

### 10x Visium spatial sequencing and analysis

The Visium Spatial Transcriptomics experiments were conducted by Singulomics Corporation (https://singulomics.com, Bronx NY). Human skin Formalin-fixed, paraffin-embedded (FFPE) tissues were sectioned onto glass slides. Deparaffinization, Hematoxylin & Eosin (H&E) staining, imaging, and decrosslinking were conducted on the tissue slides containing the FFPE tissue sections following the manufacturer’s protocol CG000520 (10x Genomics).

Following the preparation of the tissue slides, we employed the Visium assay using the CytAssist technology according to the manufacturer’s protocol CG000495 (10x Genomics). Human whole transcriptome probe panels, consisting of ∼3 pairs of specific probes for each targeted gene were added to the tissue, enabling hybridization and ligation of each probe pair. Subsequently, tissue slides and Visium CytAssist Spatial Gene Expression slides were loaded into the Visium CytAssist instrument. Gene expression probes were then released from the tissue, enabling capture by the spatially barcoded oligonucleotides present on the Visium slide surface. During this transfer process, CytAssist imaged the tissue within the fiducial frame of the Visium slide, enabling the co-registration of H&E tissue images with Visium Spatial Transcriptomics spots. Following transfer, the probes were extended and amplified and made into libraries before sequencing (paired-end 150 base reads) on an Illumina NovaSeq instrument. Space Ranger 2.0.0 (10x Genomics) was used to align CytAssist images with corresponding H&E stained images, perform quality control, and convert the Visium Spatial Transcriptomics data into spots x genes expression matrices for downstream analysis. Similar to the quality control approach used above, cells that expressed less than 500 genes, had greater than 25% mitochondrial reads, or greater than 25% of genes representing hemoglobin, were removed from the dataset. SCTransform(69) was used for sample normalization. Otherwise, the analysis pipeline was analogous to that described above using Seurat.

### In-situ hybridization

RNA in situ hybridization was performed using the RNA Scope kit (Bio-Techne, Cat # 323100) following the manufacturer’s instructions. Formalin-fixed unstained sections were first deparaffinized, treated with hydrogen peroxide blocking step, and then with protease prior to using the RNA Scope target antigen retravel. Probe hybridization and amplification steps were followed per manufacturer protocol. Slides were stained with 4’,6-diamidino-2phenylindole (DAPI) to visualize nuclei and mounted on slides using ProLong Gold Antifade (ThermoFisher, cat # P36934). *GREM1* (cat # 312831) and *CXCL10* (cat # 311851) probes were purchased from Bio-Techne. Opal 690 (Akoya Biosciences, cat # FP1497001KT) and 570 Reagent (cat # FP1488001KT) were used to visualize staining using a Stellaris 8 confocal microscope (Leica).

### Statistics

All statistical analyses were performed using the R statistical programming environment (version 4.2.3) or GraphPad Prism 10.0.2. For specific analyses, please refer to figure legends. In Figure 5, correlations were calculated using scLink v1.0.1 and corrplot v0.92 in R.

### Study Approval

The study of this cohort of patients and retrospective analysis of skin biopsies were approved by Yale Human Research Protection Program Institutional Review Boards (protocol ID 2000027571 and 2000022585, respectively). Informed consent was obtained for all patients included in single cell RNA sequencing studies. Patients and controls were recruited from Yale Dermatology clinics. Patients with CLE were biopsied on lesional skin as well as non-lesional skin of the lower back for single cell RNA sequencing. Healthy control patients were biopsied on the lower back. All biopsies performed were 4-5 mm punch biopsies. For additional patient details see **Supplementary Table 1**.

### Data Availability

All sequencing data associated with this manuscript are uploaded to NCBI Gene Expression Omnibus and will be publicly accessible as of the date 3/1/2025.

