## supplementary data for "Cutaneous lupus features specialized stromal niches and altered retroelement expression"

| Patient number | Assay | Patient group | Sex | Age | Skin Type | CLE type | Site | SLE/CLE medications |
|---|---|---|---|---|---|---|---|---|
| 1 | Single cell | CLE (+ SLE) | F | 27 | 5 | DLE | Arm | HCQ 200 mg BID + Belimumab 200 mg/ml weekly |
| 2 | Single cell | CLE | F | 26 | 2 | DLE | Arm | HCQ 200 mg BID |
| 3 | Single cell | CLE | F | 43 | 5 | DLE | Face | None |
| 4 | Single cell | CLE | F | 47 | 5 | DLE | N/A | HCQ 200 mg BID + MMF 2500 daily |
| 5 | Single cell | CLE | F | 42 | 5 | DLE | Back | Topical Clobetasol 0.05% ointment, MTX 15 mg weekly, ILK |
| 6 | Single cell | HC | F | 29 | 3 | N/A | Back | N/A |
| 7 | Single cell | HC | M | 40 | 2 | N/A | Back | N/A |
| 8 | Single cell | HC | F | 34 | 5 | N/A | Back | N/A |
| 9A | Spatial | CLE (+SLE) | F | 43 | 3 | SCLE | Arm | Pre: HCQ 200 BID + Prednisone 5 daily + Lenalidomide 5 mg qod |
| 9B | Spatial | | | | | | | Post: HCQ 200 BID + Prednisone 5 daily + Anifrolumab 300 mg q4w |
| 10 | Spatial | CLE | F | 53 | 5 | DLE | Thigh | None |
| 11 | Spatial | HC | F | 39 | 2 | N/A | Arm | N/A |

**Supplementary Table 1.** Table of patients enrolled in study including basic demographics and disease-specific medications. BID = twice daily, DLE = discoid lupus erythematosus, SCLE = subacute cutaneous lupus, HCQ = hydroxychloroquine, ILK = intralesional kenalog, MMF = mycophenolate mofetil, MTX = methotrexate, qod = every other day.

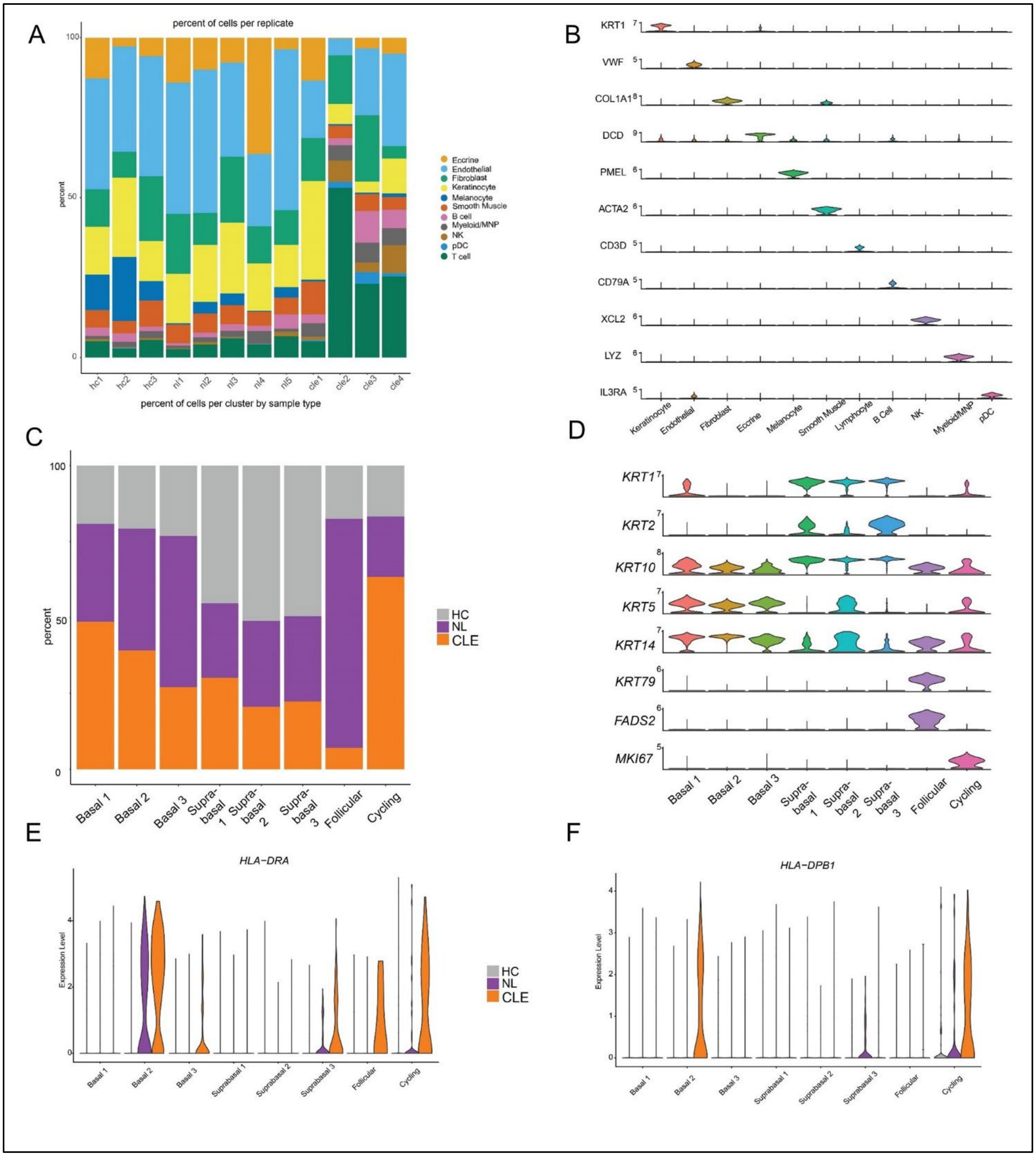


**Supplementary Figure 1.** A) Bar plot showing distribution of cell types by sample. B) Violin plot showing unique marker gene expression segregated by cell types. C) Bar plot showing distribution of cells per cluster, annotated by tissue state. D) Stacked violin plot showing unique marker gene expression of different keratinocyte populations. F and G) Violin plots of MHC II genes showing unique elevations in CLE keratinocytes.

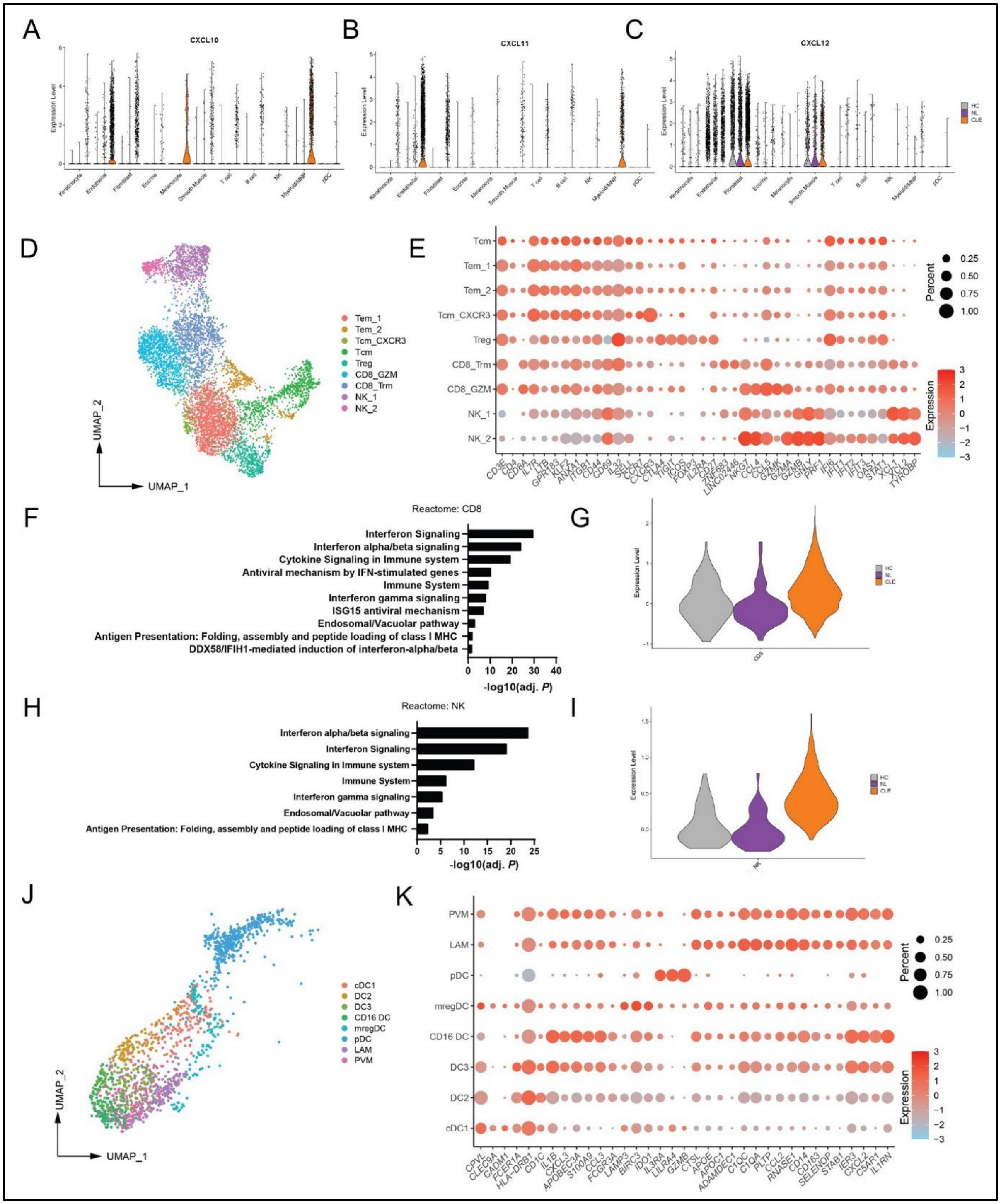


**Supplementary Figure 2.** A, B and C) Violin plots of indicated chemokines annotated by cell type and tissue cell state. D) UMAP of Lymphoid cell subclustering. E) Bubble plot of lymphoid cell marker gene expression displayed by cell type. Color indicates relative expression levels between cell types in dataset and the size indicates the percentage of cells per cluster expressing the particular gene. F and H) Reactome pathway analysis of upregulated genes in CD8 T cells (F) and NK cells (H) of CLE cells compared with HC. G and I) Violin plot of CD8 T cell cytotoxicity score and NK cell activation scores (see methods). J) UMAP plot of MNP cell subclustering. K) Bubble plot of MNP cell marker gene expression displayed by cell type.

### Canonical

| Gene | pDC | MNP |
|---|---|---|
| GZMB | ■ | |
| GPR183 | ■ | |
| PLD4 | ■ | |
| RGS2 | ■ | |
| IRF7 | ■ | |
| IRF4 | ■ | |
| IRF8 | ■ | |
| PLAC8 | ■ | |
| LILRA4 | ■ | |
| CD83 | ■ | |
| LYZ | | ■ |
| IFI30 | | ■ |
| C1QC | | ■ |
| IL1B | | ■ |
| TYROBP | | ■ |
| C1QA | | ■ |
| C1QB | | ■ |
| AIF1 | | ■ |
| HLA-DRA | | ■ |
| FCER1G | | ■ |

### Macrophage

| Gene | LAM | PVM |
|---|---|---|
| RNASE1 | ■ | ■ |
| C1QC | ■ | |
| C1QA | ■ | |
| C1QB | ■ | |
| SELENOP | ■ | |
| ADAMDEC1 | ■ | |
| PLTP | ■ | |
| APOE | ■ | |
| CD14 | ■ | ■ |
| CCL8 | ■ | |
| CXCL2 | | ■ |
| IER3 | | ■ |
| CTSL | | ■ |
| MMP19 | | ■ |
| C5AR1 | | ■ |
| IL1RN | | ■ |
| SGK1 | | ■ |
| CD163 | | ■ |

### Dendritic cell

| Gene | DC2 | DC3 | CD16 | cDC1 | mreg |
|---|---|---|---|---|---|
| CD1C | ■ | | | | |
| CD1B | ■ | | | | |
| CD1A | ■ | | | | |
| FCER1A | ■ | | | | |
| CLEC10A | ■ | ■ | | | |
| HLA-DQA1 | ■ | | | | |
| HLA-DQB1 | ■ | | | ■ | |
| HLA-DPB1 | ■ | | | ■ | |
| FCGR2B | ■ | ■ | | | |
| PKIB | ■ | | | | |
| IL1B | | ■ | ■ | | |
| PLAUR | | ■ | | | |
| LYZ | | ■ | | | |
| NLRP3 | | ■ | | | |
| CXCL8 | | ■ | ■ | | |
| CXCL2 | | ■ | | | |
| CCL3 | | ■ | ■ | | |
| TNF | | ■ | | | |
| FCN1 | | | ■ | | |
| C5AR1 | | | ■ | | |
| IL1RN | | | ■ | | |
| S100A8 | | | ■ | | |
| S100A9 | | | ■ | | |
| CCL4 | | | ■ | | |
| CXCL10 | | | ■ | | |
| ACOT7 | | | | ■ | |
| DAPP1 | | | | ■ | ■ |
| BASP1 | | | | ■ | |
| CPVL | | | | ■ | |
| PPA1 | | | | ■ | |
| ALDH2 | | | | ■ | |
| HLA-DQB2 | | | | ■ | |
| S100A10 | | | | ■ | |
| LAMP3 | | | | | ■ |
| TBC1D4 | | | | | ■ |
| GPR157 | | | | | ■ |
| FSCN1 | | | | | ■ |
| CCR7 | | | | | ■ |
| IDO1 | | | | | ■ |
| KIF2A | | | | | ■ |
| LSP1 | | | | | ■ |
| RASSF4 | | | | | ■ |

**Supplementary Figure 3:** Top 10 cell marker genes for each subclustered MNP cell type. Subclustered cell types include: cDC1, DC2, DC3, CD16+ DC, mregDC, LAM (lipid associated macrophage), and PVM (perivascular macrophage).

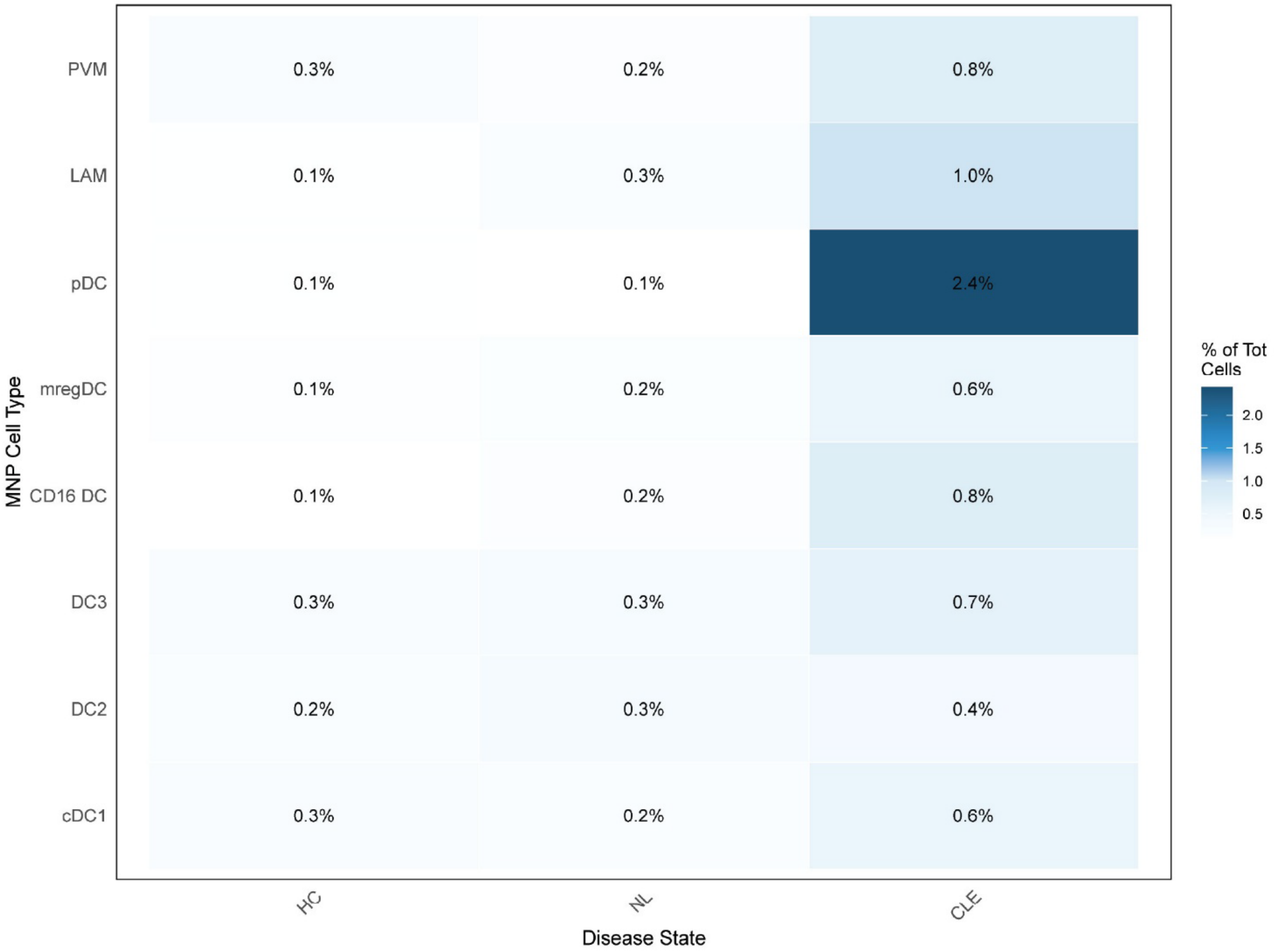


**Supplementary Figure 4:** MNP subclustered cell types, separated by tissue state. Subclustered cell types include: cDC1, DC2, DC3, CD16+ DC, mregDC, LAM (lipid associated macrophage), and PVM (perivascular macrophage).

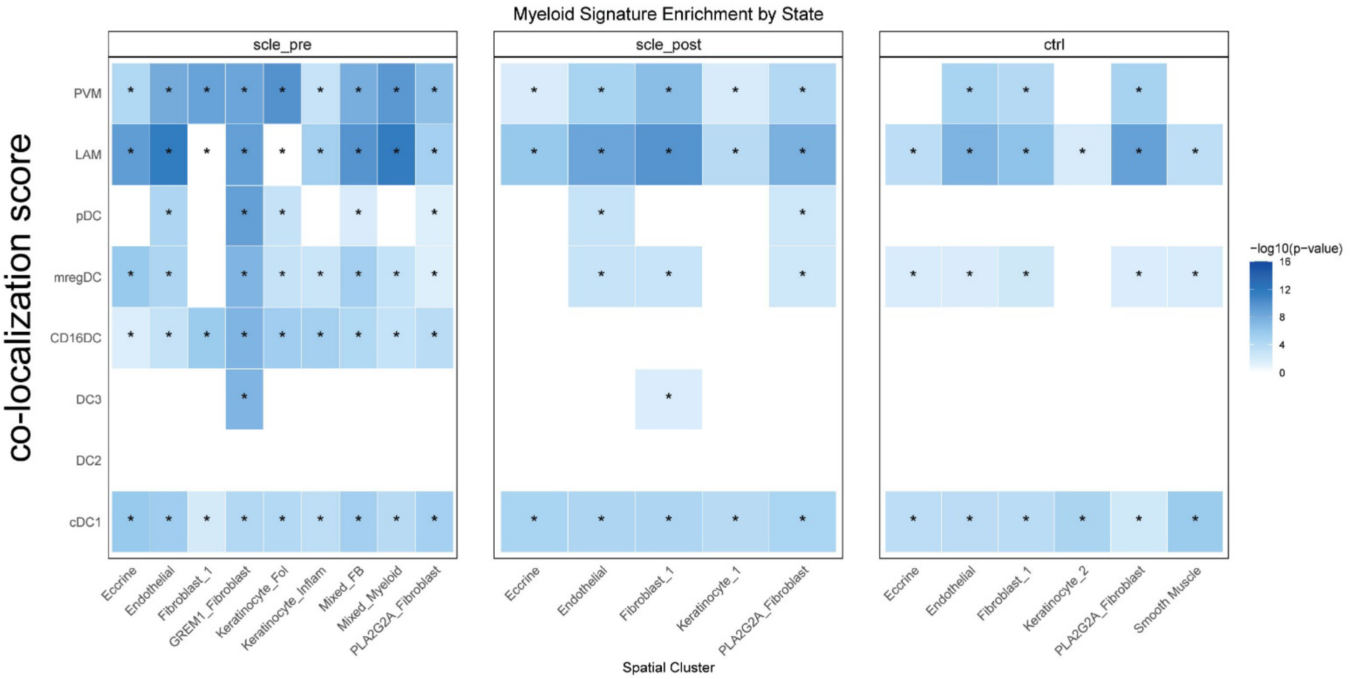


**Supplementary Figure 5:** Cell type co-localization analysis through multimodal intersection analysis, highlighting differences of Myeloid/MNP populations in different tissue states. Subclustered cell types include: cDC1, DC2, DC3, CD16+ DC, mregDC, LAM (lipid associated macrophage), and PVM (perivascular macrophage).

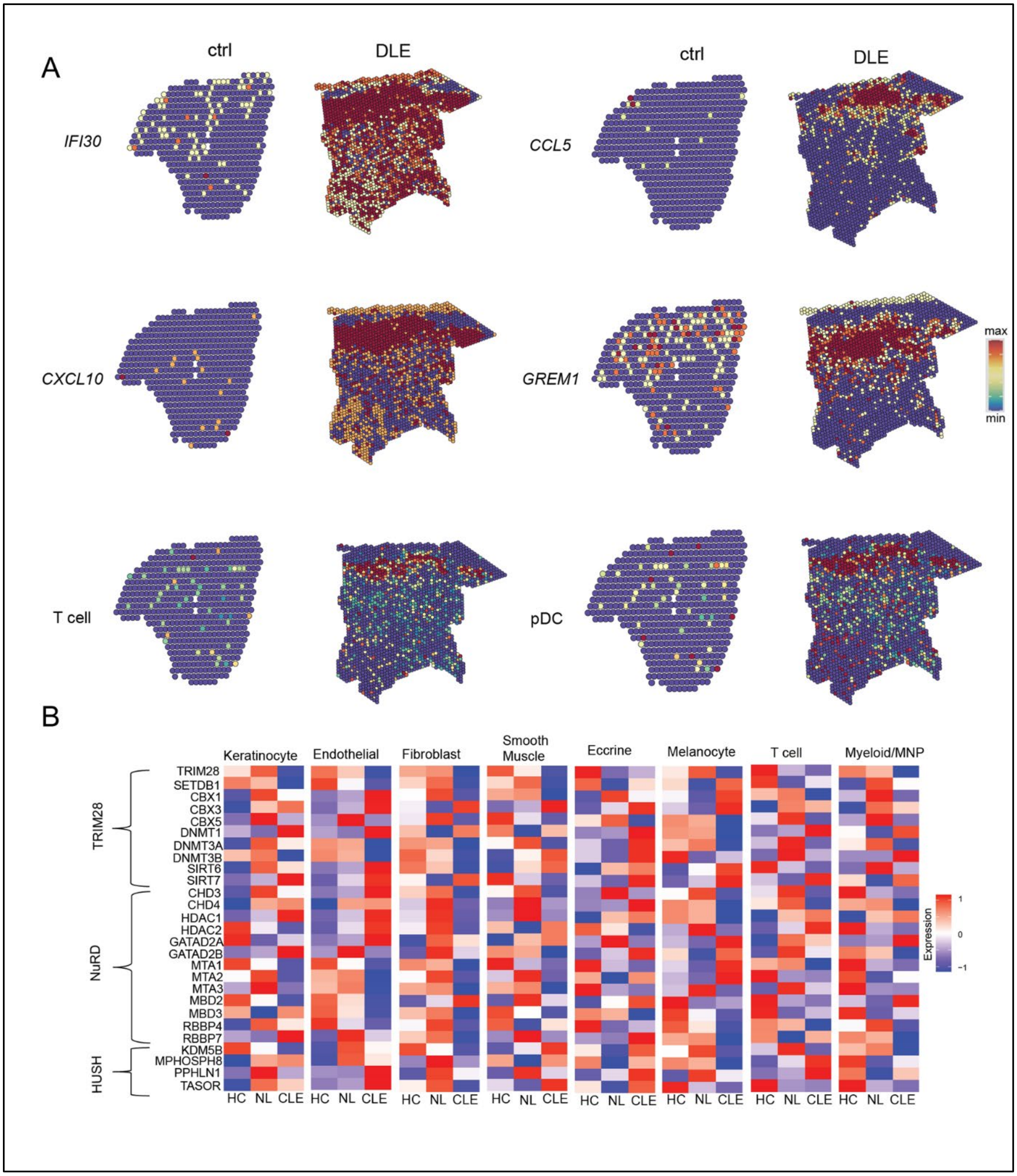


**Supplementary Figure 6.** A) Spatial gene expression plots of the indicated genes or cell type signatures (T cell, pDC) in HC (left) and DLE (right). B) Heatmap comparing expression of different epigenetic regulators segregated by tissue state and cell type.

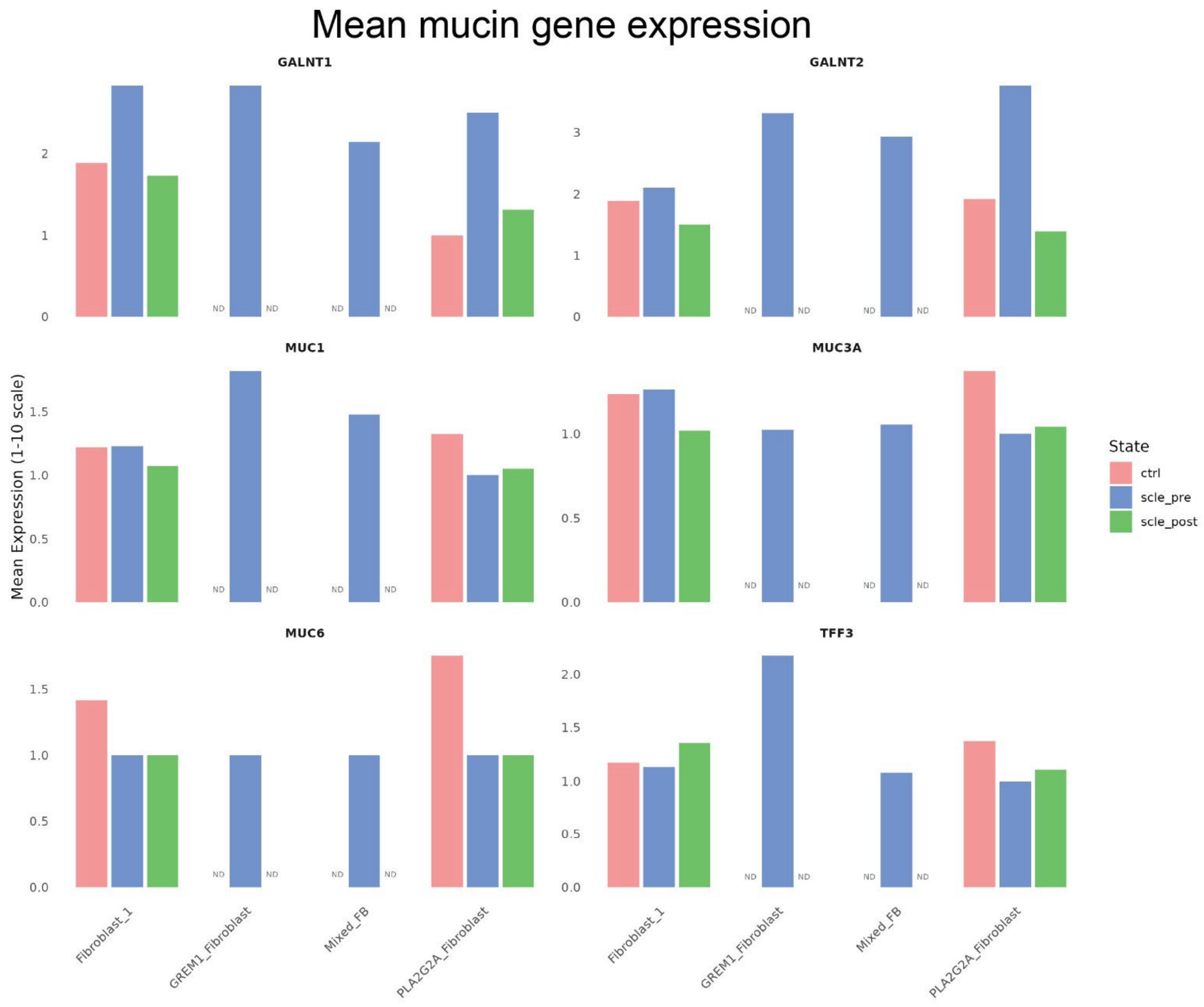


**Supplementary Figure 7.** Examination of mucin gene expression in different fibroblast clusters in pre, post-treatment, and control states. ND = cell type was not present in tissue state. ctrl = healthy control skin; scle_pre = subacute cutaneous lupus skin biopsy prior to treatment; scle_post = subacute cutaneous lupus skin biopsy post treatment with Anifrolumab (paired samples from same patient, biopsied 8 weeks a part).

A

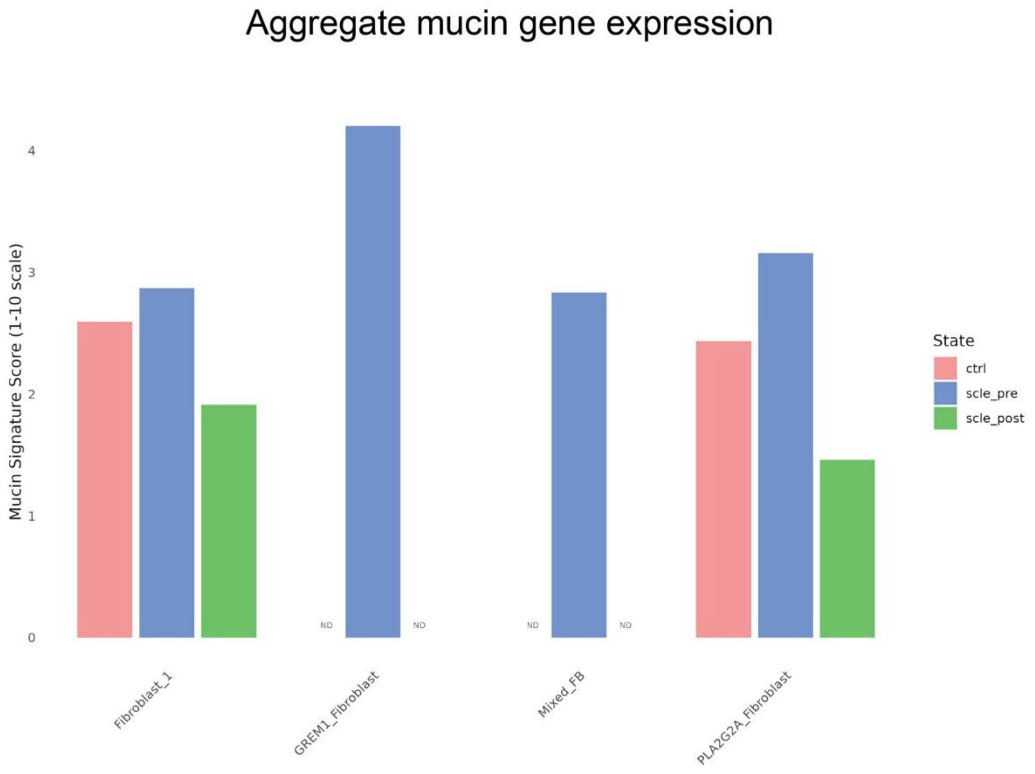


B

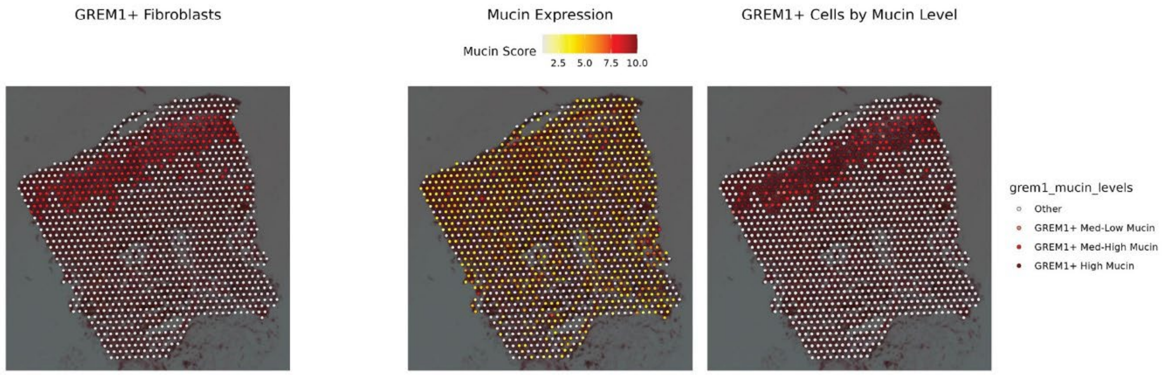


**Supplementary Figure 8.** A) Aggregate mucin gene expression (*GALNT1, GALNT2, MUC1, MUC3A, MUC6, TFF3*) in different fibroblast clusters in pre, post-treatment, and control states. ND = no cell types were present in that tissue state. B) Aggregate mucin gene expression (*GALNT1, GALNT2, MUC1, MUC3A, MUC6, TFF3*) overlaid on *GREM1*+ fibroblasts populations in CLE. The left panel plots *GREM1*+ fibroblasts and the middle panel plots the aggregate mucin gene expression. In the right panel, GREM1+ fibroblasts are plotted on the basis of low, medium, and high expression of mucin. ctrl = healthy control skin; scle_pre = subacute cutaneous lupus skin biopsy prior to treatment; scle_post = subacute cutaneous lupus skin biopsy post treatment with Anifrolumab (paired samples from same patient, biopsied 8 weeks a part).

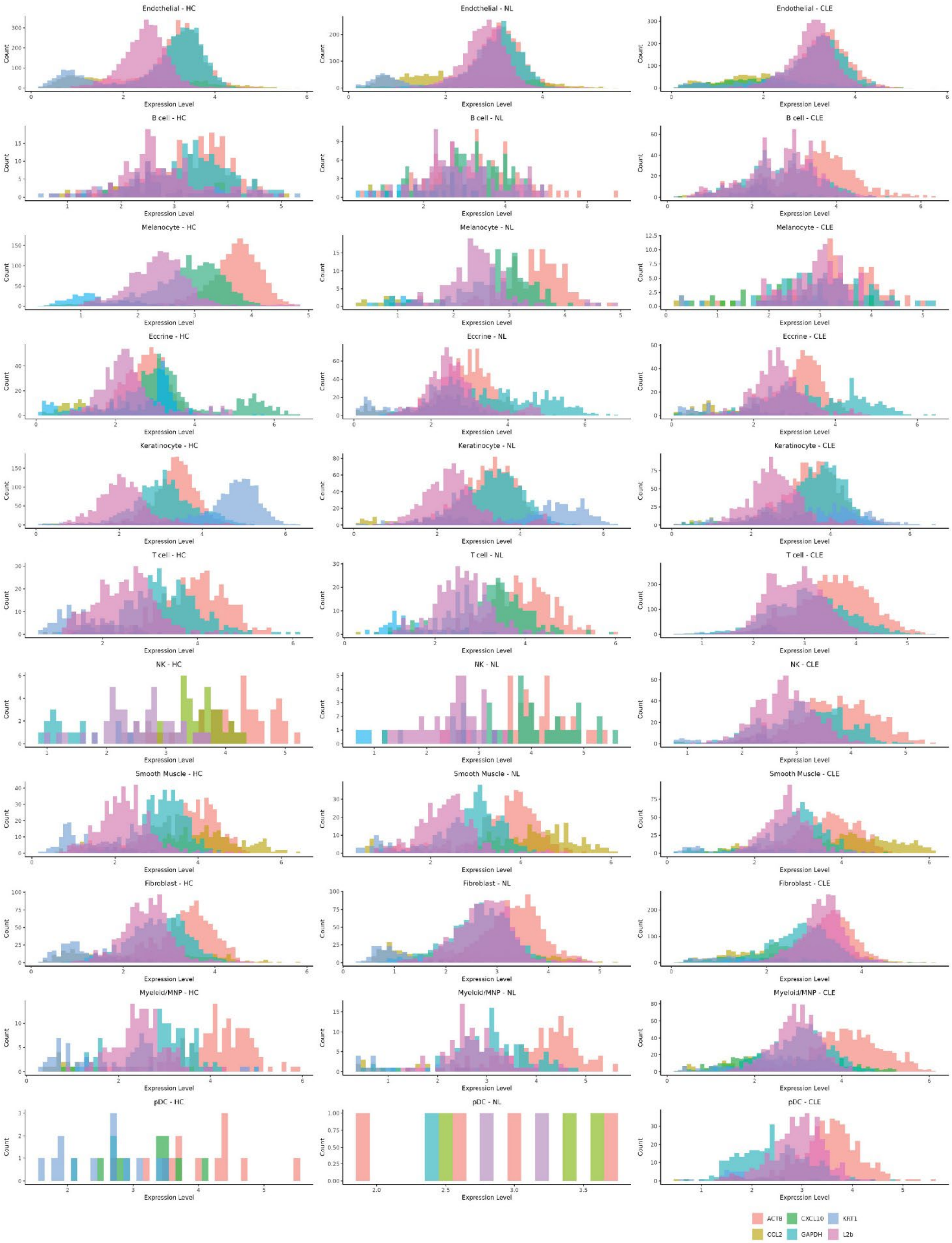


**Supplementary Figure 9.** Histogram of cell counts (y axis) and expression levels (x axis) of selected genes, including housekeeping genes (ACTB, GAPDH) as well as retroelement L2b.

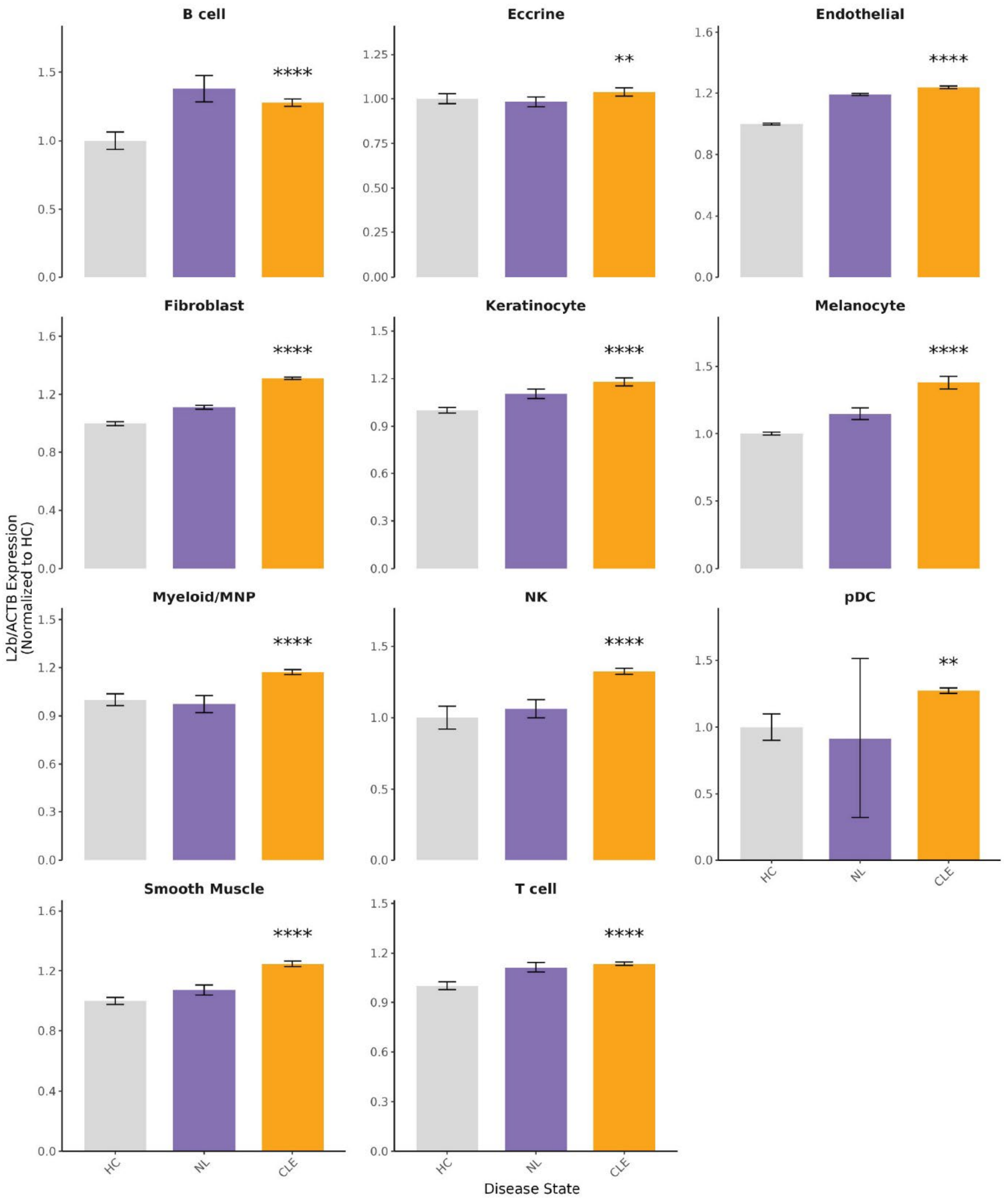


**Supplementary Figure 10.** L2b levels normalized to HC levels for all cell types and tissue states. Wilcoxon rank sum test comparing CLE and HC, * = *P* < .05, ** = *P* < .01, *** = *P* < .001, **** = *P* < .0001.

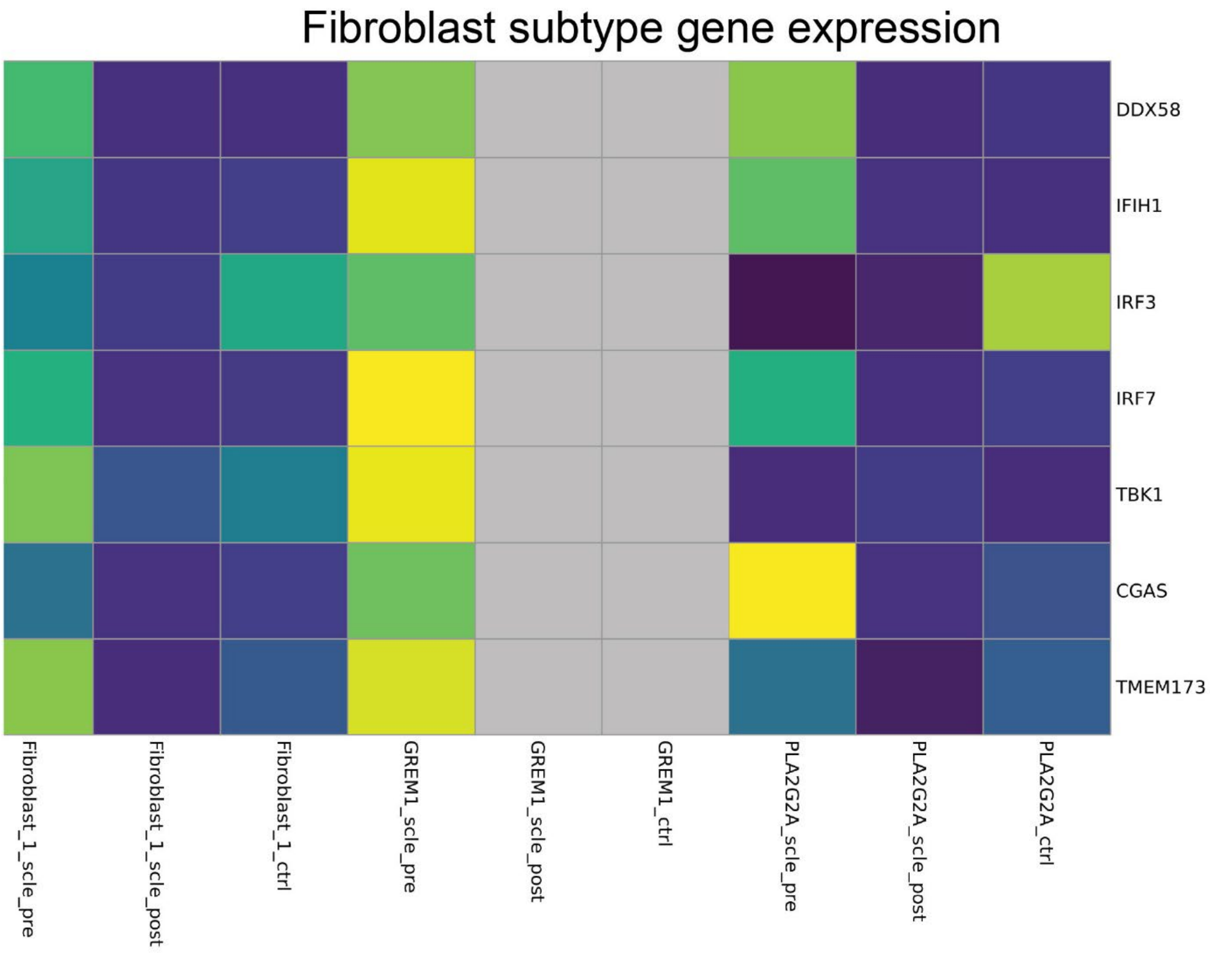


**Supplementary Figure 11.** Heatmap of RLR and cGAS-STING pathway gene expression pre and post-Anifrolumab treatment in Fibroblast clusters. These are fibroblast clusters identified in **Figure 4B**. Gray columns indicate that the fibroblast cell type is not present in that tissue state.
